# Causal Genetic Loci for a Motivated Behavior Spectrum Harbor Psychiatric Risk Genes

**DOI:** 10.1101/2023.09.06.556529

**Authors:** Jiale Xu, Romelo Casanave, Apurva S. Chitre, Qiyang Wang, Khai-Minh Nguyen, Chiara Blake, Mahendra Wagle, Riyan Cheng, Oksana Polesskaya, Abraham A. Palmer, Su Guo

## Abstract

Behavioral diversity is critical for population fitness. Individual differences in risk-taking are observed across species, but underlying genetic mechanisms and conservation are largely unknown. We examined dark avoidance in larval zebrafish, a motivated behavior reflecting an approach-avoidance conflict. Brain-wide calcium imaging revealed significant neural activity differences between approach-inclined versus avoidance-inclined individuals. We used a population of ∼6,000 to perform the first genome-wide association study (GWAS) in zebrafish, which identified 34 genomic regions harboring many genes that are involved in synaptic transmission and human psychiatric diseases. We used CRISPR to study several causal genes: serotonin receptor-1b (*htr1b*), nitric oxide synthase-1 (*nos1*), and stress-induced phosphoprotein-1 (*stip1*). We further identified 52 conserved elements containing 66 GWAS significant variants. One encoded an exonic regulatory element that influenced tissue-specific *nos1* expression. Together, these findings reveal new genetic loci and establish a powerful, scalable animal system to probe mechanisms underlying motivation, a critical dimension of psychiatric diseases.

---

Antipredation, attention deficit and hyperactivity disorder (ADHD), depression, EKW zebrafish, exploration, genetic influences of natural behavioral variation, genetics of quantitative traits, genome editing, risk-taking, schizophrenia.

## Introduction

The nervous system produces behavioral responses that are influenced by genetics, the environment, and their interactions ^1-5^. Behavioral variation is commonly observed among individuals of the same species. Studies of behavioral variation in genetic model organisms have provided insights into sensory, motor, drug consumption, and decision-making related processes ^6-12^, but the extent of conservation of underlying genes and pathways is largely unknown.

Individual differences in risk-taking versus safety-seeking have been observed across species. In humans, excessive risk-taking or safety-seeking is associated with psychiatric diseases ^13,14^. For instance, individuals with attention deficit and hyperactivity disorder (ADHD) or bipolar disorder (in the manic phase) are more likely to engage in risky behaviors, whereas individuals with anxiety disorders have unreasonable and overwhelming fears that drive them to seek safety. Genome-wide association studies (GWAS) in humans have implicated genetic loci in psychiatric diseases. However, it remains challenging to identify the underlying causal genes and mutations and to understand their function in broad biological contexts ^15^.

Light as a sensory stimulus affects cognition and mood ^16-18^. Light-dark preference is a behavior that is observable across species ^19-23^ and considered an anxiety-like trait in mammals ^24^. Larval zebrafish generally avoid dark that mimics a predator shadow, and this dark avoidance behavior, enhanced by stressors and alleviated by commonly used anxiolytics ^25,26^, is naturally variable in the population and such variability is heritable ^27-29^.

Larval zebrafish as young as 5 days post fertilization (dpf) are free-living and need to hunt for food and avoid predators, thus functional neural circuitry for exploration and antipredation is in place. Zebrafish larvae are available in large numbers, amenable to high throughput behavioral phenotyping and brain-wide neural activity imaging and circuit analysis ^30^. At a young age, their behavior is minimally affected by cumulative environmental effects, making larval zebrafish a more sensitive system than adults for detecting genetic influences.

In this study, we showed that dark avoidance in larval zebrafish reflected a classical approach-avoidance conflict ^31^: wanting to approach dark for exploratory foraging (taking risks), versus wanting to avoid dark for antipredation (seeking safety). Brain-wide neural activity imaging uncovered that the behavioral spectrum of approach-avoidance was underscored by activity differences in distinct brain regions. We performed the first GWAS using 5,759 larval zebrafish to identify causal genes and variants that drive this motivated behavior. We discovered new genetic loci, and furthermore causal genes, which showed strong conservation from fish to men, with many being associated with psychiatric diseases, most prominently, schizophrenia, depression, and attention deficit and hyperactivity disorder (ADHD).

## Results

### Variation of a Motivated Behavior Underscored by Neural Activity Differences

#### Light-dark preference: a behavioral readout for approach-avoidance conflict

We hypothesized that variation of dark avoidance in larval zebrafish reflected an individualized strategy for resolving a motivational conflict between approach versus avoidance. We tested 6-7 dpf larval zebrafish from an outbred EKW (EKkwill Wildtype) line using a multi-trial light-dark preference behavioral paradigm ^27-29^. The light-dark choice index (LDCI), which computes % time in dark - % time in light, provides a quantitative measure of the behavior. Individuals with LDCI (averaged across 4 trials) in the top 90% are termed variable dark avoidance (VDA), as their dark avoidance behavior tends to be variable across trials, whereas individuals with LDCI in the bottom 10% are termed strong dark avoidance (SDA), as they spend hardly any time in the dark side (**Figure 1A**). In addition to the LDCI, SDA and VDA groups also differed in Total Dark zone Entry Number (TDEN), Average Dark zone Entry Duration (ADED), Latency to First Dark zone Entry (LFDE), and Total Distance traveled in the Dark Zone (TDDZ), whereas the velocity and Total Distance traveled (TD) were not different (**Figure S1A**).

**Figure 1.**
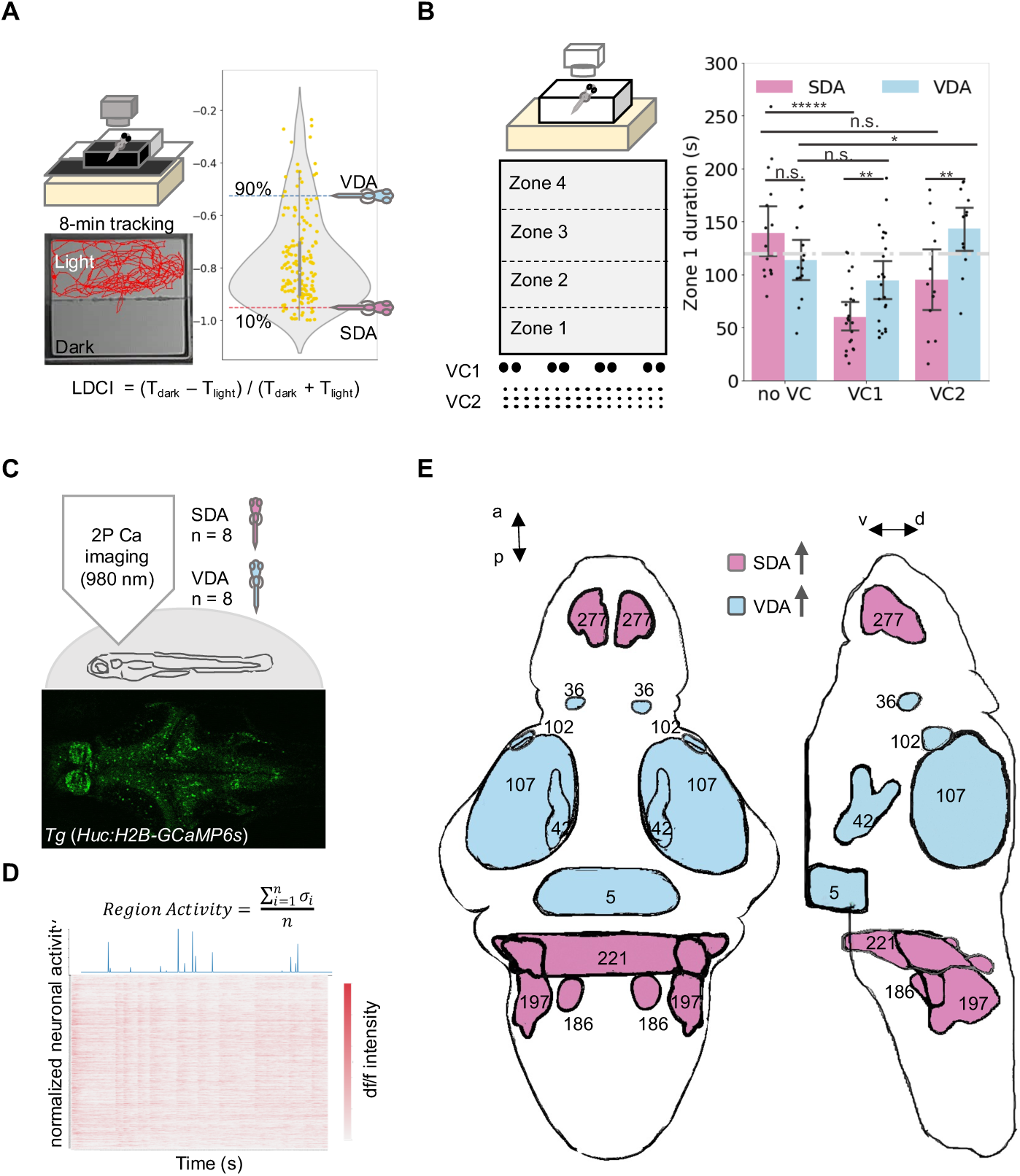
Individual variation in light-dark preference reflects differences in motivation toward food-or predator-mimicking cues and is underscored by brain activity differences. **A**. Individual variation of light-dark preference in a larval population (n=192) quantified by the Light-Dark Choice Index (LDCI). Individuals with LDCI (averaged across 4 trials) above the 90^th^ percentile were grouped as variable dark avoidance (VDA) while those with LDCI below the 10^th^ percentile were grouped as strong dark avoidance (SDA). **B**. Behavioral differences of approaching food-mimicking versus predator-mimicking visual cues (VC) in SDAs and VDAs. Two-sample t-test, n=12-23 per group per visual cue condition, \**p*<0.05, \*\**p*<0.01, \*\*\*\*\**p*<0.00001, n.s. not significant. Mean ± s.e.m. **C.** *In vivo* brain-wide calcium imaging of spontaneous neural activity in SDAs and VDAs (8 dpf) (n=8 for each group). See also **Movie S1**. **D**. Quantification of brain activity at cellular resolution based on GcAMP6s signal (dF/F) time series (900 s, at 1Hz). Mean variance across neurons assigned to an anatomical mask is computed as the average activity of the mask. **E.** Anatomical masks with significant differences of neuronal activity between SDAs and VDAs. In comparison with VDA, SDA shows hyperactivity in four masks (red; 277. Telencephalon - Isl1 cluster 1; 221. Rhombencephalon - Rhombomere 3; 186. Rhombencephalon - Medial Vestibular Nucleus; 192. Rhombencephalon - Neuropil Region 4); and hypoactivity in five masks (blue; 36. Diencephalon - Isl1 cluster 1; 102. Mesencephalon - Retinal Arborization Field 7 (AF7); 107. Mesencephalon - Tecum Neuropil; 42. Diencephalon - Migrated Posterior Tubercular Area (M2); 5. Diencephalon - Caudal Hypothalamus). a-p, anterior-posterior; v-d, ventral-dorsal. One-way ANOVA / ANCOVA with permutation test (see methods).

Individuals belonging to the VDA and SDA groups were then selected and subjected to a new behavioral test at 9 dpf, using visual cues that mimic food (for stimulating exploratory foraging) or predator eye spots (for evoking antipredation). This test consisted of four trials of 8-min tracking in an open tank arena, with one of two sets of visual cues (VC) presented on one side wall of the arena: VC1 consisted of 4 pairs of black dots that resemble eye spots of predators (diameter = 2 mm), whereas VC2 consisted of 48 small black dots that occupy approximately the same area as eye spots and mimic food particles (diameter = 0.5 mm). Larval zebrafish have been previously shown to avoid large (predator-like) and to approach small (food-like) particles ^32,33^. We also examined a third condition in which no visual cues were presented as a control (see **Figure 1B**). For data analysis, the open tank is divided into four equal virtual zones, with Zone 1 closest to the VCs. In the absence of VCs, both SDAs and VDAs spent about equal time in Zone 1, averaging ∼120 seconds (about ¼ of the total time in the arena). In the presence of predator mimicking VC1, SDAs spent significantly less time in Zone 1. In the presence of food mimicking VC2, VDAs spent significantly more time in Zone 1, but SDAs did not (**Figure 1B**). Together, these results suggest that individual variation in dark avoidance reflects a classical approach-avoidance conflict: VDAs are approach-inclined (i.e., hyper-exploratory, risk-taking), SDAs are avoidance-inclined (i.e., hyper-anti-predatory, safety-seeking), with the rest of the population somewhere in between.

#### Brain-wide neural activity differs between approach-inclined VDAs and avoidance-inclined SDAs

To uncover potential neural basis underlying the behavioral differences between VDA and SDA groups, we performed brain-wide calcium imaging of spontaneous neuronal activity, a measure that likely reflects functional wiring and internal brain state. Following phenotyping of *Tg[HuC-H2BGCaMP6s]* larval zebrafish that express the genetically encoded and nuclear localized calcium indicator H2B-GCaMP6s in neurons, individual SDAs or VDAs were identified and subjected to whole brain calcium imaging under a 2-photon microscope (**Figure 1C, Figure S1,** and **Movie S1**). The fluorescence of calcium indicators was converted using the CalmAn software ^34^ to obtain individual neuronal activity traces. Registration to the Z-brain atlas ^35^ allowed us to calculate average neuronal activity in each anatomical mask across individuals (**Figure 1D, Figure S1B-C**). These analyses revealed that SDAs displayed higher activity in more rostral (e.g., #277, telencephalic Islet cluster 1) and caudal (e.g., #211, Rhombencephalon Oxtl Cluster 1 Sparse) brain regions, whereas VDAs displayed higher neuronal activity in diencephalic regions including the migrated posterior tubercular area (M2) (#42), the caudal hypothalamus (#5), and Islet1 cluster (#36) (**Figure 1E, Figure S1D**). Thus, individual behavioral differences are underscored by neuronal activity differences in distinct brain regions, with VDAs displaying higher activity in the diencephalic brain regions including the hypothalamus.

### Discovery of GWAS Loci Associated with the Motivated Behavior Variation

#### GWAS workflow

Individual variation of motivation-associated light-dark preference in larval zebrafish is a quantitative trait. To uncover genes and pathways controlling this behavioral variation, we carried out GWAS using larval zebrafish from the outbred EKW population (**Figure 2A**). Pairwise crosses with rotated mating among 162 male and 142 female breeders generated a population of 6,216 larval zebrafish, of which 6,054 had successfully recorded behavioral data. Computational analysis of video recordings produced quantitative behavioral phenotype data. To remove environmental co-variates, we considered factors that accounted for >1% of the variance. These included the effects of arenas, batches, and groups. Quantile normalization was subsequently performed to address any non-normally distributed phenotypes.

**Figure 2.**
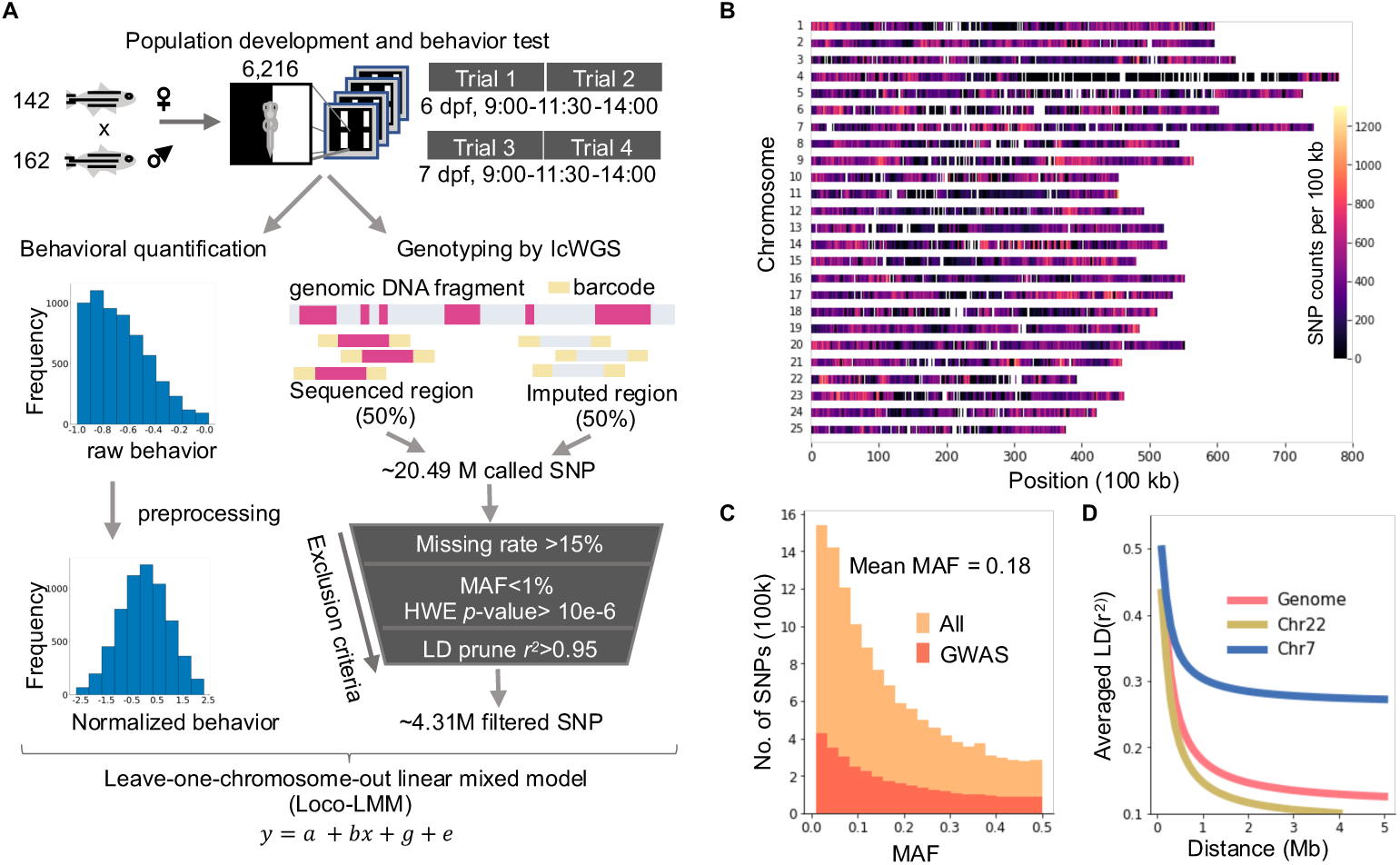
Genome-wide association study (GWAS) employing ∼6K phenotyped and genotyped larval zebrafish. **A**. Workflow of GWAS. All genotyping data were deposited to SRA (ID pending). HWE, Hardy-Weinberg equilibrium; lcWGS, low coverage whole-genome sequencing; LD, linkage disequilibrium; SNP, single-nucleotide polymorphism. **B**. Genome-wide distribution map of LD-pruned 4.35 million SNPs in the EKW population. Density per 100 kb was color-coded and shown for all 25 chromosomes. **C**. MAF (Minor Allele Frequency) distribution is shown for the 20.49 million imputed SNPs (orange) and for the 4.3 million SNPs used for GWAS (red). Mean MAF is the same (0.18) for both sets. **D**. Plot showing the LD (Linkage Disequilibrium) decay using averaged r^2^ for SNPs spaced up to 5 Mb apart. The LD decay across the genome was fitted in a quadratic curve that reaches below 0.2 within 1 Mb (red). LD distribution fitted for individual chromosomes shows fastest LD decay in chr 22 and slowest in chr 7.

Genomic DNA was prepared following individual behavioral phenotyping. A two-step procedure as previously described in mice ^36^ was used for genotyping. We first performed a low-coverage whole genome sequencing (lcWGS) to obtain reads randomly distributed across the genome. We then used the whole population to call variants by imputation. To perform imputation, we constructed a reference panel from the 94 deep-sequenced parental breeders. The sequences generated for all subjects were deposited to SRA (ID pending). Our genotyping pipeline yielded more than 20 million polymorphic SNPs, nearly double the mean of three other zebrafish strains ^37^. After quality control and linkage disequilibrium (LD) pruning, 4.31 million SNPs were retained for GWAS analysis.

#### Genetic diversity and a *rapid decay of linkage disequilibrium (LD) are observed in the zebrafish GWAS population*

We used imputed genotypes to investigate population structure and cryptic relatedness that might be confounding for GWAS (**Figure S2**). The samples did not cluster after projection onto the first 4 principal components, suggesting that there is no obvious population stratification. Individual inbreeding coefficient estimated by *plink* displayed no correlation with PC1, and its distribution showed no inbreeding in our samples.

The distribution of 4.31 million SNPs on all 25 chromosomes showed a non-uniform distribution (**Figure 2B**). In particular, the large segment of low SNP density on chromosome 4 corresponds to its long arm region with a lack of protein-coding genes and an extensive heterochromatin state ^38^.

One population genetic feature that determines the GWAS power is minor allele frequency (MAF). An assessment based on all SNPs showed that most SNPs (78%) had MAF greater than 0.01. The MAF of the LD-pruned SNPs had an average MAF of 0.18 (**Figure 2C**). To estimate the mapping resolution, we calculated the decay rate of LD using SNPs separated up to 5 Mb (**Figure 2D**). A more rapid LD decay was found in our population in comparison to mammalian populations ^36,39^, highlighting a strength of the zebrafish EKW population for high-resolution genetic mapping.

#### GWAS loci for the motivated behavior

We analyzed seven quantitative behavioral indices as shown in **Figure S1A**. Four are light-dark preference related: Light-Dark Choice Index, Total Dark zone Entry Number, Average Dark zone Entry Duration, and Latency to First Dark zone Entry. Three are related to swimming velocity or distance traveled: Velocity, Total Distance traveled, and Total Distance traveled in Dark Zone. Each index is composed of five sub-indices for parameters derived from each trial (T1 to T4) and their mean (**Figure 3A**). Using SNP-based inheritance measures ^40^, we detected considerable heritability (**Figure S3A**) and correlation among these behavioral indices at both phenotypic and genotypic levels (**Figure S3B**).

**Figure 3.**
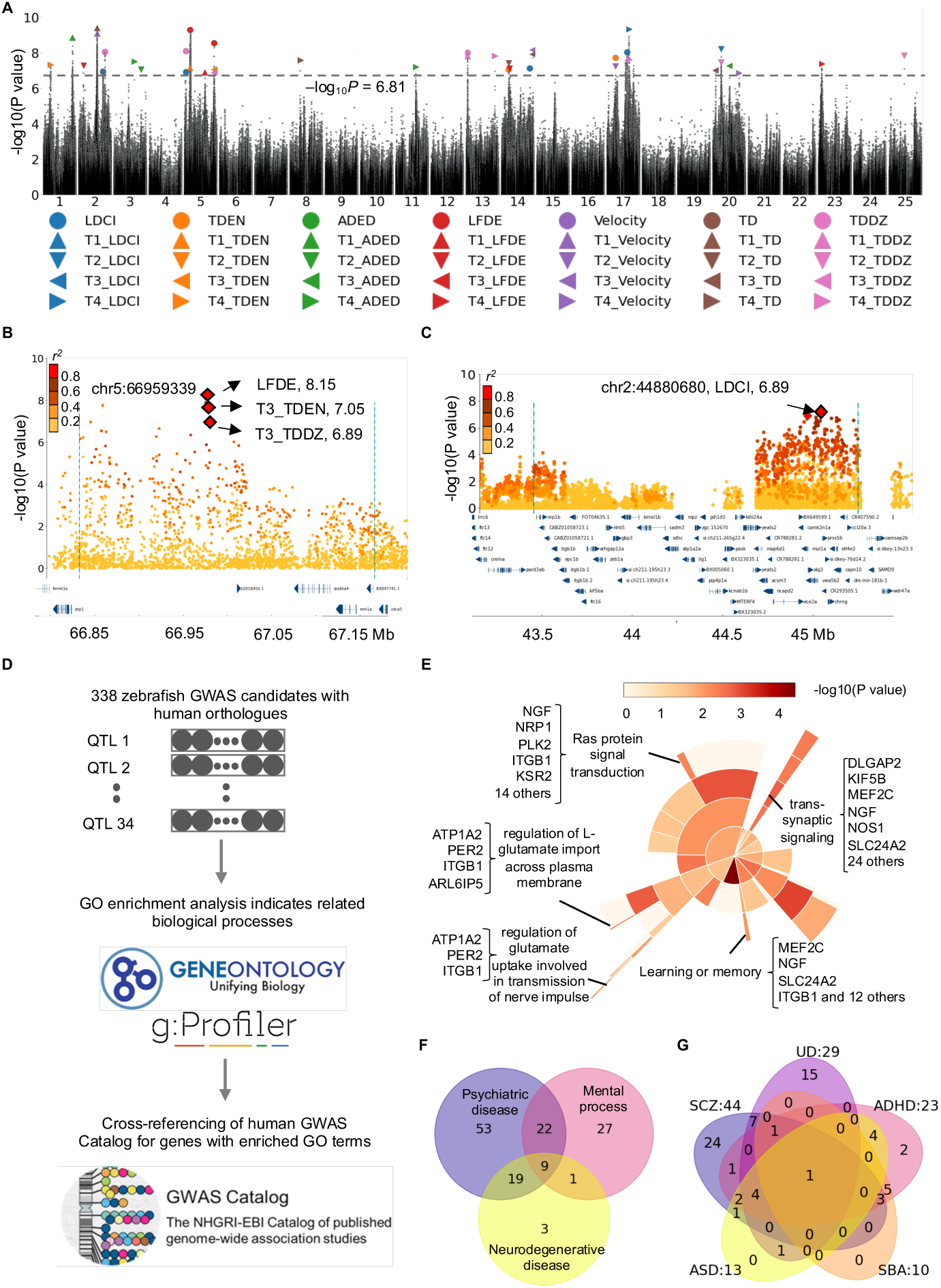
GWAS reveals causal genetic loci with candidate genes involved in synaptic transmission and human psychiatric diseases. **A.** Manhattan plot shows the identified 34 QTLs for 35 behavioral indices. The top SNPs for each behavioral index were marked. The black line indicates a permutation-derived significance threshold of –log_10_*P* = 6.81 (*p* = 1.4 x 10^-7^). LDCI (Light-Dark Choice Index), TDEN (Total Dark zone Entry Number), ADED (Average Dark zone Entry Duration), LFDE (Latency to First Dark zone Entry), TD (Total Distance traveled), and TDDZ (Total Distance traveled in the Dark Zone). **B-C.** Example Locus Zoom plots. (**B**) A pleiotropic locus at chr 5 (66.8-67.1 Mb) identified for three behavioral indices with the same top SNP. (**C**) An example QTL (chr 5, 30.6 – 31.5 Mb) with many candidate genes. **D**. Schematic of the approach linking zebrafish GWAS candidate protein-coding genes to functional pathways and human GWAS findings. 338 candidate genes with human orthologues are cross-referenced with Gene Ontology, gProfiler, and human GWAS catalog database. **E**. Sunburst plot of molecular process terms enriched in human orthologues of the zebrafish GWAS candidate genes. Each piece is one term, and the degree of the redness indicates enrichment significance. **F.** Venn diagram shows the number of human orthologues in categories “psychiatric disease”, “mental process” and “neurodegenerative disease”. **G.** Venn diagram partitions the 102 psychiatric disease-related genes into five categories. ADHD: attention deficit hyperactivity disorder; ASD: autism spectrum disorder; SBA: substance abuse; SCZ: schizophrenia; UD: unipolar depression.

To identify the associated loci, we applied the leave-one-chromosome-out Linear Mixed Model (LOCO-LMM) ^41^. We performed 1,200 rounds of permutation to derive the significance threshold of −log_10_(*P*-value) = 6.81 (*P* = 1.4 x 10^-7^) (**Figure S3C**). Using this threshold, we identified 40 significant loci (**Figure 3A**). For some behavioral indices, the loci were mapped nearby one another on the same chromosome. To determine whether they support multiple independent QTLs, we used stepwise regression to calculate correlations between the top SNP and other neighboring SNPs. Correlated SNPs were then combined and represented by the top SNPs. This resulted in a total of 34 GWAS loci.

To determine the genomic regions for each GWAS locus, we identified markers that had a high correlation coefficient with the top SNPs (r^2^>0.6). The resulting 34 genomic regions ranged from ∼7 kb to ∼4 Mb in size. Representative Locus Zoom plots for two GWAS QTLs were shown (**Figure 3B-3C**).

### GWAS Candidate Genes Are Involved in Synaptic Transmission and Human Psychiatric Diseases

We identified 338 candidate protein-coding genes that had human orthologues (**Figure 3D**). The Gene Ontology (GO) enrichment analyses of these genes uncovered significantly enriched biological processes including trans-synaptic signaling, glutamatergic neurotransmission, RAS protein signal transduction, and learning or memory (**Figure 3E**).

By cross-referencing the GWAS candidate genes with enriched GO terms to the human GWAS Catalog ^42^, we noted that more genes were in the categories of psychiatric diseases and mental processes versus neurodegenerative diseases (**Figure 3F**), with 53 psychiatric disease-unique vs. 3 neurodegenerative disease-unique genes. Among the 102 genes associated with human psychiatric diseases, more genes was observed for schizophrenia (24 category-unique), unipolar depression (15 category-unique), and attention deficit hyperactivity disorder (ADHD) (2 category-unique), versus autism spectrum disorders (ASD) (0 category-unique) and substance abuse (SBA) (0 category-unique) (**Figure 3G**).

### Causal genes for motivated behavior regulation: *htr1b*, *stip1,* and *nos1*

To prioritize candidate genes for causality validation, we ranked top SNPs based on their *p-*values and selected small LD blocks containing relatively few candidate genes. We further prioritized genes based on their likely expression in the nervous system, links to significant GO terms, and association with human GWAS findings (**Figure 4A**). Control genes were also included from both within and outside the LD blocks.

**Figure 4.**
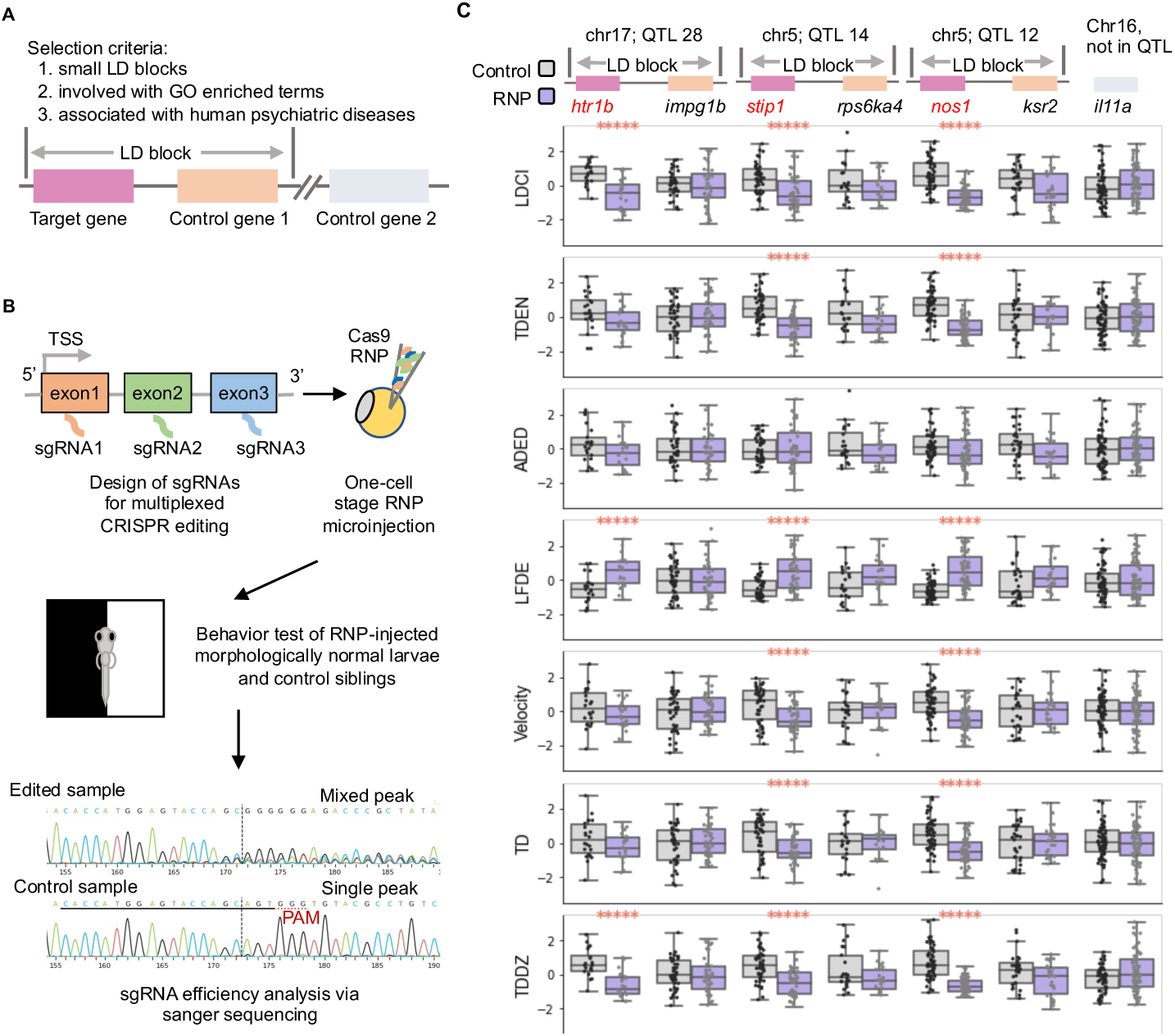
Multiplexed CRISPR genome editing uncovers causal genes for the motivated behavior. **A**. Gene selection criteria for causality test. At least one control gene from the same QTL region is tested together (orange) with the target gene (purple). One control gene, *il11a*, located outside of all the QTLs is used as a universal control gene (gray). **B**. Schematic of the multiplexed *Crispr-Cas9* mediated gene knock-out (KO) pipeline. **C**. Comparisons of seven behavioral parameters between RNP-injected groups and the sibling controls. Of the seven tested genes, three from 3 different QTLs were validated as causal genes. Two-sample t-test*, p*<0.05, \*\**p*<0.01, \*\*\**p*<0.001, \*\*\*\**p*<0.0001, \*\*\*\*\**p*<0.00001. n.s. not significant, n=22-90. For each gene, Ns are the same between control and RNP-injected. *p*-values are adjusted by *Benjamini–Hochberg* false discovery rate correction for test across all indices. *htr1b (serotonin receptor 1b)*, *impg1b (interphotoreceptor matrix proteoglycan 1b), stip1 (stress-induced phosphoprotein 1)*, *rps6ka4 (ribosomal protein S6 kinase a4), nos1 (neuronal nitric oxide synthase 1), ksr2* (*kinase suppressor of ras 2*), *il11a* (*interleukin 11a*).

For instance, in the QTL #28 (708 kb) on Chromosome 17 with the highest peak SNP’s *-logP* value of 9.3 (**Figure S4A**), three protein-coding genes were selected from a total of six for functional characterization: *serotonin receptor 1b (htr1b)*, *interphotoreceptor matrix proteoglycan 1b (impg1b)*, and *myosin VI b (myo6b)*. We also evaluated another small QTL #14 (171 kb) on Chromosome 5 with the top SNP’s *-logP* value of 8.2 (**Figure S4B**), two out of five protein-coding genes were selected for functional study: *stress-induced phosphoprotein 1 (stip1)*, and *ribosomal protein S6 kinase a4 (rps6ka4)*. We next examined a large LD block that contained many genes: the QTL #12 (1.3 Mb) on Chromosome 5 with the top SNP’s *-logP* value of 8.7 and 33 protein-coding genes (**Figure S4C**). The gene *neuronal nitric oxide synthase 1 (nos1)* is expressed in the central nervous system, is part of the GO term “trans-synaptic signaling”, and is associated with multiple psychiatric disorders including schizophrenia, unipolar depression, insomnia, and post-traumatic stress disorders. We therefore selected *nos1*, as well as a nearby gene, *kinase suppressor of ras 2* (*ksr2*). The gene *interleukin 11a* (*il11a*), which is not located in any of our QTLs, was also selected as a control.

To functionally test these genes, we established a scalable CRISPR knockout (KO) pipeline in G_0_ animals injected with the Cas9 ribonucleoprotein (RNP) complexes. Three computationally predicted high efficacy sgRNAs were selected and synthesized for each candidate gene; RNP complexes of Cas9 and 3 sgRNAs were micro-injected into one-cell stage zebrafish embryos, followed by behavioral testing at 6-7 dpf and subsequent genotyping for sgRNA KO efficiency (**Figure 4B**). We used the EKW breeders that give rise to larval progeny with an average level of dark avoidance in the medium range, to enable observation of KO-induced increases or decreases in dark avoidance.

As a control, the delivery of Cas9 protein alone did not affect any of the 7 behavioral indices (**Figure S4D**). *In vivo* efficacy of the sgRNAs used to target these genes were evaluated following behavioral phenotyping. The efficacies for inducing both Indels (insertions and deletions) and KOs (knockouts of the proteins resulting from frameshifts and stop codons) ranged from 40% to 100% (**Figure S4E**). We also evaluated the three sgRNA efficacy prediction algorithms by correlating the predicted efficacies with the *in vivo* observed efficacies. Among them, CRISPRscan gave the best correlation (r=0.48) whereas IDT showed the lowest correlation (r=0.09) (**Figure S4F**).

For QTL #28, we found that knocking out *myo6b* resulted in morphological defects and were therefore not used in subsequent behavioral studies. Knocking out *htr1b* but not *impg1b* significantly altered the behavior, with decreased light-dark choice index (indicative of decreased time spent in the dark zone), increased latency to first dark zone entry, and decreased dark zone total distance traveled. The total dark zone entry number and average dark zone entry duration showed a decreasing trend, whereas the velocity and total distance traveled were not significantly affected. For QTL #14, we found that knocking out *stip1* but not *rps6ka4* significantly altered all behavioral indices except the average dark zone entry duration. For QTL #12, we found that knocking out *nos1* but not *ksr2* significantly altered all behavioral indices except the average dark zone entry duration. Additionally, knocking out the *il11a* had no significant effects on any of the behavioral indices under study (**Figure 4C**).

In summary, we found that KO of *htr1b*, *stip1, and nos1*, which were in three QTL regions, increased the motivation-associated dark avoidance behavior in larval zebrafish.

### Genome-wide Analysis of GWAS Significant Variants Uncovers Those with Evolutionary Conservation and Predicted Regulatory Function

Our GWAS was run on 4.31 million LD pruned SNPs, as there is little gain to use all 20.49 million discovered SNPs but a considerable increase in computational costs. To uncover putative causal variants that account for the motivated behavior variation in the population, we used the unpruned SNP set to perform association mapping for each QTL region. This resulted in 1,288 significant SNPs, which were then analyzed using three approaches (**Figure 5A**). First, we used the Ensembl Variant Effect Predictor to classify these variants into predicted functional categories. Second, we determined the conservation of sequences harboring these significant SNPs across 8 vertebrate species that are composed of five fish species (fugu, medaka, stickleback, and tetraodon in addition to zebrafish), frogs (x. tropicalis), mice, and humans. This analysis uncovered 52 conserved elements harboring 66 GWAS significant variants, sized from 11 to 2,298 base-pairs with conservation scores ranged from 0.38 to 0.76. Third, we used the DANIO-CODE database ^43^ to uncover 40 zebrafish putative regulatory elements that contain at least one GWAS significant variant.

**Figure 5.**
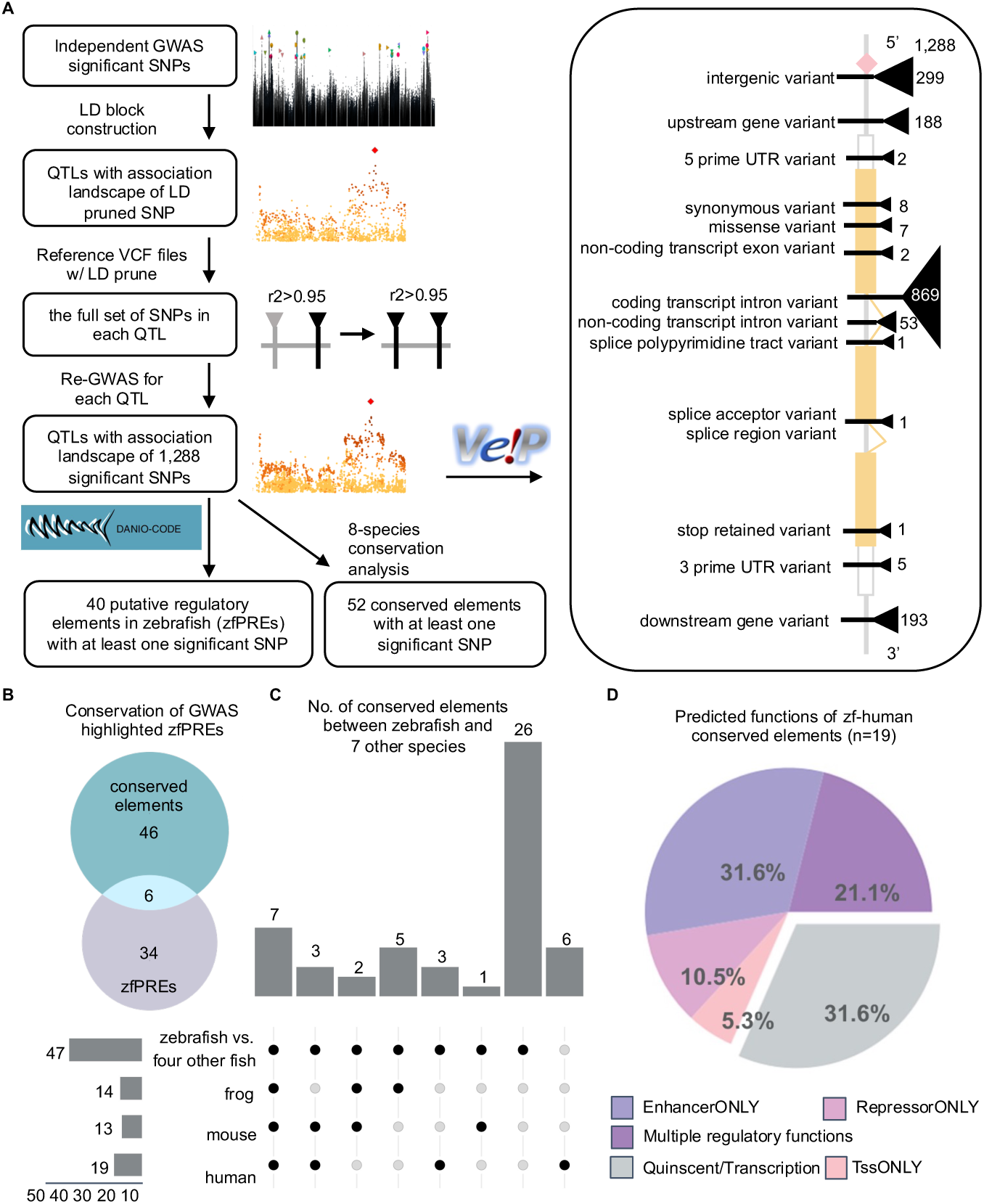
Genome-wide analysis of GWAS significant variants. **A**. Pipeline for QTL-based Re-GWAS to uncover all significant variants and bioinformatics analyses using VEP (Variant Effect Prediction), DANIO-CODE database, and UCSC 8-species conservation datasets. **B**. Overlap between the conserved elements and zebrafish putative regulatory elements (zfPRE). **C**. Upset plot of the conserved elements between zebrafish and 7 other species. The bar charts show number of conserved elements in each conservation category (vertical) and each species (horizontal). **D**. Functional categories of human conserved elements, predicted by the RegulomeDB.

Among the 52 conserved elements, 6 were putative zebrafish regulatory elements predicted by the DANIO-CODE database (**Figure 5B**). Analysis of the 52 conserved elements further uncovered that 7 were conserved across all 8 vertebrate species, and 19 were conserved between zebrafish and humans (**Figure 5C**).

Since the regulatory landscape of the zebrafish genome is not as well defined compared to the human genome, we leveraged the conservation and the availability of regulatory information in the mammalian genomes to predict the function of the 19 conserved elements found in humans. Using the RegulomeDB v2.2 ^44^, we found that most elements have predicted regulatory function, either assuming multiple functions, or as enhancers only or repressors only. About one third of the elements are predicted to be quiescent or transcribed (**Figure 5D**). Together, these analyses identify the GWAS significant and putative causal variants and further accentuate those that are in evolutionary regions and have predicted regulatory functions.

### An Evolutionarily Conserved Exonic Regulatory Element Containing a Putative Causal Variant is Required for Proper Spatial Expression of *nos1*

We next focused on a DANIO-CODE predicted cis-regulatory element that contains a GWAS significant variant. This element is conserved across all 8 vertebrate species examined. It is located 234 kb upstream of the *nos1* gene and encodes an exon of the neighboring *ksr2* gene (**Figure 6A**).

**Figure 6.**
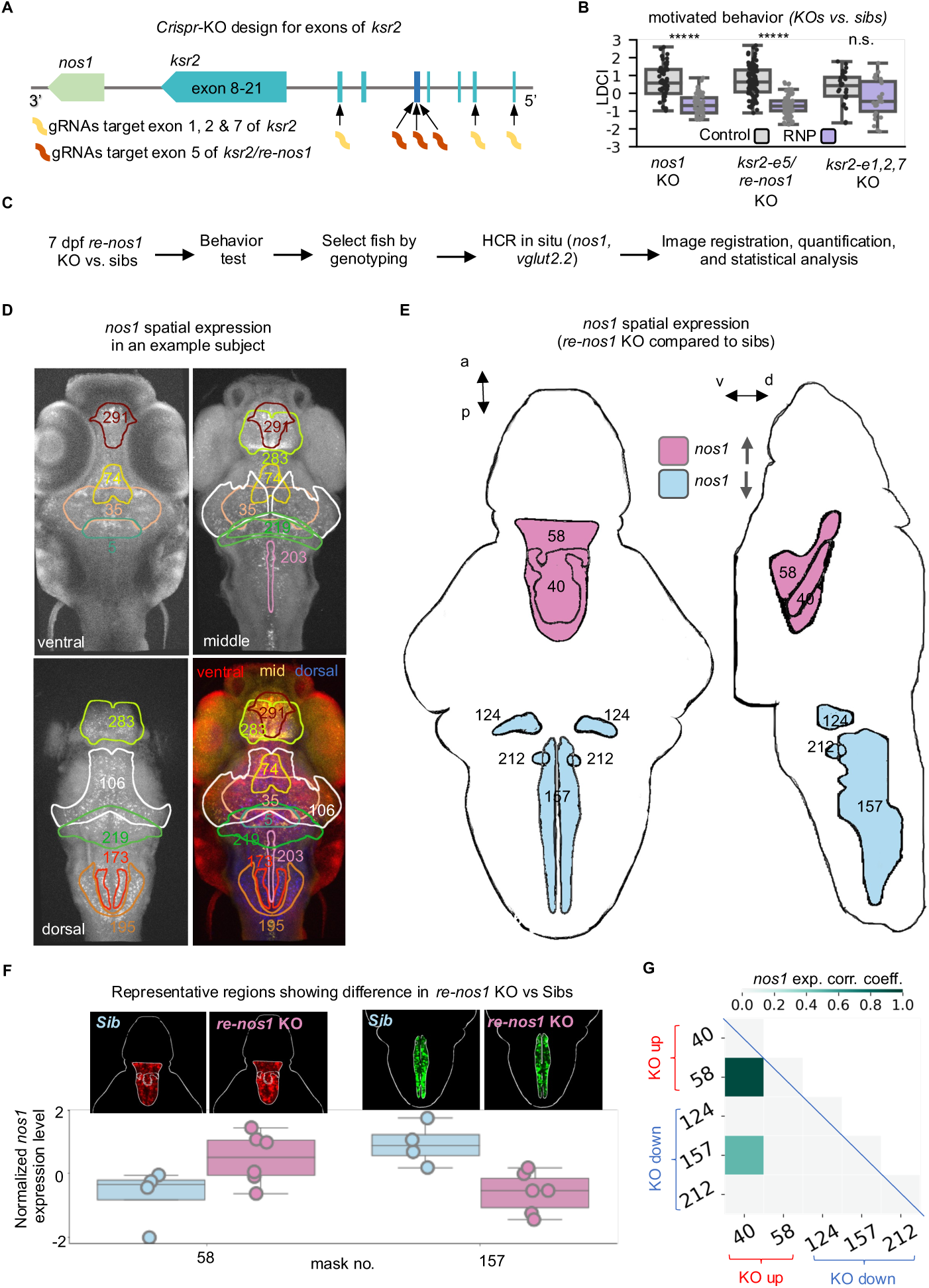
Discovery of an exonic regulatory element that is required for the motivated behavior and *nos1* spatial expression. **A**. Schematic of multiplexed genome editing targeting *ksr2* exons. The exon is numbered based on the largest transcript of *ksr2*. All targeted exons are shared across most *ksr2* transcripts. The exon 5 (*ksr2-e5*, dark blue), 234 kb upstream of *nos1,* is predicted to have enhancer activity in the DANIO-CODE database, hence also named *re-nos1* (putative regulatory element for *nos1*). **B**. LDCI (Light-Dark Choice Index) in *nos1* KO, *ksr2-e5/re-nos1* KO, and ksr2-e1,2,7 KO. Two sample t-test, \*\*\*\*\**p* < 0.00001, n.s. not significant., n=27-90, Ns are the same between control and RNP-injected. *Benjamini–Hochberg* FDR control across all indices. **C**. Flow chart showing the analysis of *nos1* spatial expression across the whole brain at cellular resolution using HCR *in situ*. The *nos1* HCR is multiplexed with *vglu2.2* for imaging data registration. **D**. Example fish with *nos1* expression in the whole brain. Related z-planes were projected to show *nos1* expression in ventral, middle, dorsal, and the whole brain. Representative masks for *nos1* expression are labeled and numbered based on the Z-brain atlas. See also **Movie S2. E**. Differential expression of *nos1* comparing *re-nos1* KO to sibling controls. Two Diencephalon masks (red) were detected with upregulated *nos1* expression in the *re-nos1* KO. Three Rhombencephalon regions (blue) showed downregulated *nos1* expression in the *re-nos1* KO. **F**. Example brain regions with increased (left) and decreased (right) *nos1* expression in the *re-nos1* KO compared to sibling controls. **G.** *Pearson* correlation of anatomical masks with altered *nos1* expression in the *re-nos1* KO vs sibling controls.

To characterize the function of this putative exonic regulatory element, we designed sgRNAs to target this exon (*ksr2-e5/re-nos1*) as well as other exons of *ksr2 (ksr2-e1,2,7)*. CRISPR KO of this, but not other *ksr2* exons, significantly increased dark avoidance, with the behavioral indices showing remarkable similarity to the KO of *nos1* gene (**Figure 6B**, and **Figure S5**). These results suggest that it is not the *ksr2* exonic role of this element that is critical for the motivated behavior, but rather, it may be its regulatory role on *nos1* that is critical for the behavior.

To further explore this idea, we performed spatial gene expression analysis of *nos1*, using the Hybridization Chain Reaction (HCR) *in situ* technology ^45,46^ on *re-nos1* KO and siblings (**Figure 6C**). The *re-nos1* KO animals carried ∼300 bp deletion in the *re-nos1* element that included the GWAS significant variant (**Figure S5**). Gene-specific probes were designed for *nos1* and *vglut2.2*, the latter of which was used for brain registration to obtain anatomical information and as a control (**Figure S6**). We carried out quantitative image analysis, which uncovered that in 7 dpf larval zebrafish, the time at which motivated behavior was assessed, *nos1* expression was detected in distinct brain cell types distributed from the telencephalon to the rhombencephalon, including the pallium, sub-pallium, hypothalamus, and various small anatomical masks located within these regions (**Figure 6D**). 232 Z-brain masks expressed both *nos1* and *vglut2.2*. Among 53,824 (232 x 232) *nos1*-*vglut2.2* expression correlations analyzed, a small subset of positive (1.4%, n=760) and negative (0.05%, n=32) correlations showed significance, suggesting that the expression of these genes is mostly but not entirely independent. We also analyzed intra-correlation of 257 Z-brain anatomical masks that expressed *nos1* or *vglut2.2*. For *nos1*, 5,201/32,896 (15.8%) correlations are significantly positive whereas 302/32,896 (0.9%) correlations are significantly negative. For *vglut2.2*, 4,068/32,896 (12.4%) correlations are significantly positive whereas 1,197/32,896 (3.6%) correlations are significantly negative.

Compared to siblings, *re-nos1* KO animals had significantly increased *nos1* expression in two diencephalic areas: Posterior Tuberculum (#58) and Medial vglut2 cluster (#40), and significantly decreased *nos1* expression in three hindbrain areas: Anterior Cluster of nV Trigeminal Motor neurons (#124), Gad1b strip 3 (#157), and Oxtl Cluster 2 Near MC axon cap (#212) (**Figure 6E**). Two representative regions with altered *nos1* expression in *res-nos1* KOs were shown (**Figure 6F**). Cross correlation analysis uncovered that the two *nos1*-up masks (#58 and # 40) were significantly correlated with each other while *nos1*-down masks were not (**Figure 6G**). We also analyzed *vglut2.2* expression and found three regions showing differences between *re-nos1* KO and siblings; *nos1* expression in these regions were not altered, and *vglut2.2* expression in these regions was not correlated with *nos1* expression.

Among the anatomical regions with altered *nos1* expression, it is worth noting that the posterior tuberculum is anatomically homologous to the human substantia nigra and contains dopaminergic neurons that project to the sub-pallium (homologous to the mammalian striatum) ^47^. These striatal pathways control both motivation and initiation of movement ^48^. Together, these results reveal that an evolutionarily conserved element harboring a GWAS significant variant is required to regulate the motivation-associated dark avoidance behavior, likely due to its role in tissue-specific regulation of *nos1* expression.

### Selection of Approach-inclined versus Avoidance-inclined Traits Significantly Impacts Tissue-specific Expression of *nos1*

To understand whether altered *nos1* expression may partially underlie natural variation of motivated behavior, we performed spatial gene expression analysis on SDA and VDA individuals, which were selectively bred for five generations (**Figure 7A**). The impact of selective breeding on the motivated behavior was evident, as the light-dark choice index and other related behavioral indices showed a clear trend of separation across generations (**Figure 7B**) (**Figure S7**). The starting population had a behavior somewhat skewed toward dark avoidance. As a result, we observed a weaker selection effect on SDA than VDA.

**Figure 7.**
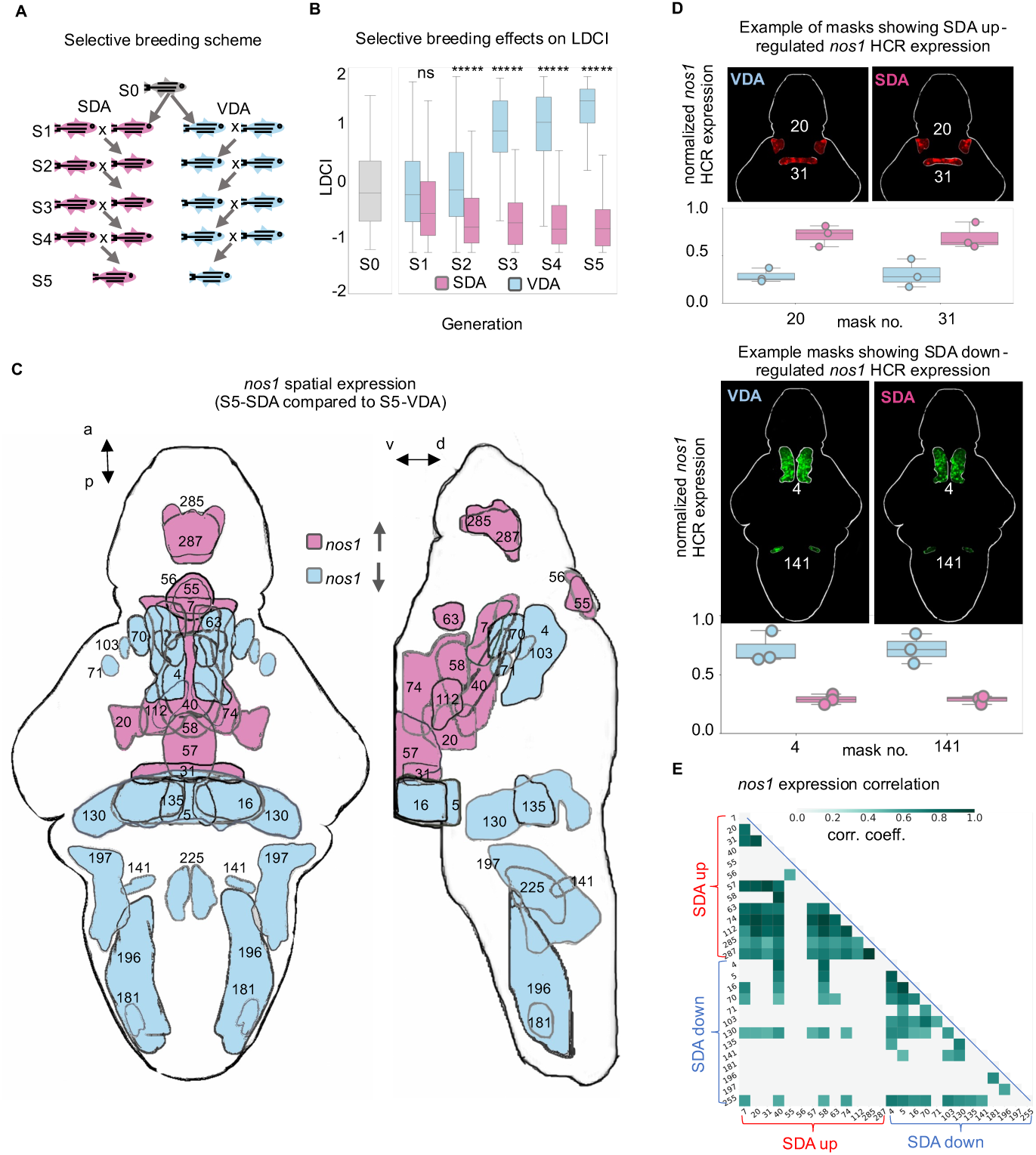
Differential *nos1* expression in approach-inclined VDA and avoidance-inclined SDA individuals. **A**. Schematic of selective breeding for SDA and VDA over five generations. The founder (gray) is a subset of the GWAS population. Multiple S1 families are developed and selected fish are always crossed with fish from different families to avoid inbreeding effects. **B**. Accumulation of selective breeding effects on LDCI (Light-Dark Choice Index) across generations. Two-sample t-test, \*\*\*\*\**p*<0.00001, n.s. not significant., n=35 for each generation. *p*-values are adjusted by *Benjamini–Hochberg* false discovery rate correction for test across all indices. **C**. Comparison of *nos1* spatial expression in the larval zebrafish brain between the 5^th^ generation of SDAs and VDAs. The S5-SDA differs from S5-VDA with increased *nos1* expression in 13 brain regions and decreased *nos1* expression in 13 other brain regions. Most of the detected diencephalon regions show increased *nos1* expression in S5-SDA whereas the Rhombencephalon regions show decreased *nos1* expression. **D**. Example masks with SDA up-regulated (top, red) and down-regulated (bottom, green) *nos1* expression. **E**. *Pearson* correlation of anatomical masks with altered *nos1* expression.

HCR *in situ* for *nos1* and *vglut2.2* was performed on the 5^th^ generation SDAs and VDAs, following behavioral phenotyping. We uncovered region-specific differences of *nos1* expression between SDAs and VDAs, in overlapping but broader domains than the differences observed in *re-nos1* KO animals compared to siblings (**Figure 7C**). Compared to VDAs, SDAs had significantly increased *nos1* expression in medial regions of the diencephalon, including the Posterior Tuberculum (#58) and Medial vglut2 cluster (#40); both regions also showed increased *nos1* expression in *re-nos1* KO animals compared to siblings. SDAs had decreased *nos1* expression in several other regions of the diencephalon [e.g., anterior pretectum cluster of vmat2 neurons (#4), retinal arborization field 4 (#70)], and regions in the mesencephalon and rhombencephalon. Two representative regions with altered *nos1* expression in an SDA versus a VDA were shown (**Figure 7D**). Cross correlation analysis uncovered that *nos1*-up masks are more corelated with one another than *nos1*-down masks (**Figure 7E**). We also analyzed *vglut2.2* expression and found six regions showing differences (5 up and 1 down) in SDA compared to VDA groups. Most were not correlated with *nos1* expression. These results suggest that natural genetic variations in the EKW population influence the approach-avoidance spectrum in part by regulating tissue-specific expression of *nos1*.

## Discussion

We characterized a motivation-associated light-dark preference behavior at the larval zebrafish stage when environmental imprinting on behavior is low compared to adult stages and the basic neural circuitry for exploration and antipredation is already in place. We then performed the first GWAS in zebrafish, which uncovered 34 QTLs. We validated several causal genes spanning three QTLs, *htr1b*, *stip1*, and *nos1,* all of which are associated with human psychiatric diseases ^49-51^. We further found that an evolutionarily conserved sequence element harboring a putative GWAS causal variant, which also encodes an exon of a neighboring gene, is required to regulate proper *nos1* expression and this motivated behavior. Both increased and decreased *nos1* expression was observed when this element was knocked out, suggesting that it has multiple functions of activating and repressing *nos1* in a tissue-specific manner; alternatively, some of the observed effects may be indirect. The observation of altered *nos1* expression in naturally approach-inclined VDAs versus avoidance-inclined SDAs provide evidence leading us to propose that natural genetic variations influence this motivated behavior in part through tissue-specific modulation of *nos1* expression.

*Nos1* encodes a neuronal nitric oxide (NO) synthase that is a downstream signaling molecule of N-methyl-D-aspartate receptor (NMDAR) activation. Decreased NOS1 expression has been observed in the anterior cingulate cortex of depressive patients ^52^. NO acts on the cyclic guanosine monophosphate (cGMP)-dependent kinase (PKG) to regulate neuronal excitability ^50^. Intriguingly, the study of “rovers” and “sitters”, two naturally occurring variants in Drosophila food-search behavior has previously uncovered a role of PKG ^11^, which also underlies age-related transition from hive work to foraging in honeybees ^12^. Taken together, our findings provide evidence that evolutionarily conserved genes and pathways regulate motivation. They further establish larval zebrafish as a powerful *in vivo* experimental system to elucidate both the molecular and cellular basis of motivated behavior.

Individuals on opposite ends of the light-dark preference behavioral spectrum differ significantly in their motivation for approaching small food-like dots and for avoiding large predator-like eye spots, as well as their neuronal activity brain-wide. Approach-inclined VDAs displayed higher activity in diencephalic regions including the hypothalamus, a brain structure that modulates the perception of food and the willingness to take risks in foraging decisions in larval zebrafish ^33^. In contrast, avoidance-inclined SDAs exhibited higher activity in telencephalic and hindbrain regions, which are more tuned to sensorimotor activities. These results reveal that natural genetic variations “program” brain activity differences across individuals to implement behavioral variation in a population. Different alleles likely promote fitness in different environments. Future investigations of this motivated behavior, the identified QTLs, and causal genes in wild populations would allow us to further explore this idea.

The expression of psychological and behavioral traits was vividly described by Charles Darwin ^53^. Studies of genetic influences on behavior in simpler model organisms have suggested that humans are typical of other animal species ^4^. In these contexts, it is perhaps not surprising that all three causal genes we discovered thus far, *htr1b*, *nos1*, and *stip1,* have been linked to human psychiatric diseases ^49-51^. What is unexpected is that, given the seemingly different behavioral phenotypes, *that is*, the extent of dark avoidance in larval zebrafish versus clinical symptoms of psychiatric diseases, one third of our GWAS candidate genes are associated with psychiatric diseases, with preferential links to depression, schizophrenia, and ADHD. What connects fish and human behavioral phenotypes, however, is that they all reflect functional outputs of motivational circuitries, the mechanistic understanding of which remain incomplete and fragmentary. Causal validation of other candidate genes using the larval zebrafish system holds promise not only to understand fundamental principles of motivated behavior regulation but may also shed important lights on the causal genetics for human psychiatric diseases, which is a major challenge facing the research community, and has benefited from molecular genetic approaches using model organisms ^54-56^. Larval zebrafish provide a high throughput system to study the motivation-associated light-dark preference, making this system suitable for small molecule screening ^57-61^, which may yield new therapeutics to address the unmet medical needs for psychiatric diseases.

Zebrafish as a vertebrate model organism have been widely used for *in vivo* functional studies at the level of individual genes. As the first GWAS study employing zebrafish, this work has demonstrated the utility of this model organism for uncovering the causal genetics of quantitative phenotypic traits. Given the considerable conservation between zebrafish and humans at the genomic, anatomical, and physiological levels, we anticipate that future quantitative genetic studies using outbred zebrafish will not only provide fundamental insights into complex traits but also help decipher human disease mechanisms and aid in therapeutic discovery.

## ACKNOWLEDGMENTS

We thank the UCSF Imaging Core Facility for support, Michael Munchua, Ginger Huang, Lena Cook, and Sydney Gonzalez for fish care, Kevin Anderson for assistance with fish breeding, members of the Guo and Palmer labs for helpful discussions. This research was supported by NIH R01 GM132500 and DA053385 (S.G.).

## AUTHOR CONTRIBUTIONS

S.G. and A.P. conceived the project. J.X. led all aspects of the research and performed most experimental work and analysis, with assistance from R.C., Q.W., C.B., and M.W. A.S.C., K.N., R.C., and O.P. performed genotyping and GWAS data curation. J.X. and S.G. wrote the paper, with inputs from all authors.

## METHODS

### KEY RESOURCES TABLE

| REAGENT or RESOURCE | SOURCE | IDENTIFIER |
| --- | --- | --- |
| Chemicals, peptides, and recombinant proteins |  |  |
| Cas9-NLS purified protein | QB3 MacroLab, UC Berkeley | https://qb3.berkeley.edu/facility/qb3-macrolab/ |
| Mivacurium chloride | Abcam | Cat#ab143667 |
| Critical commercial assays |  |  |
| Twist 96-Plex Library Preparation kit | TWIST, San Francisco, CA | https://www.twistbioscience.com/products/ngs/library-preparation/twist-96-plex-library-preparation-kit |
| KAPA Hyper Library Preparation kit | Roche | Cat#7962363001- KK8504 |
| DNAdvance Genomic DNA Isolation Kit | Beckman Coulter | Cat#A48706 |
| HCR RNA-FISH Probe Set | Molecular Instruments | N/A |
| Deposited data |  |  |
| Reads from low-cove WGS generated by Twist 96-Plex Library Preparation kit | This paper | SRA: TBD |
| Reads from WGS generated by KAPA Hyper Library Preparation kit | This paper | SRA: TBD |
| VCF genotype data | This paper | TBD |
| Code and analyzed data | This paper | TBD |
| Experimental models: Organisms/strains |  |  |
| Zebrafish: Tg(*Huc:H2B-GCaMP6s*) | <sup>62</sup> | JF5 |
| Zebrafish: EKW | EKKWill | http://www.ekkwill.com/ |
| Oligonucleotides |  |  |
| Primers for multiplexed *Crispr-Cas9* gene knock-out | Integrated DNA Technologies | N/A |
| Guide RNAs for multiplexed *Crispr-Cas9* gene knock-out | Integrated DNA Technologies | N/A |
| Software and algorithms |  |  |
| Noldus Ethovision XT | Noldus | https://www.noldus.com/ethovision-xt |
| ImageJ 2.14.0 | <sup>63</sup> | https://imagej.nih.gov/ij/ |
| GCTA 1.93.2 | <sup>64</sup> | https://yanglab.westlake.edu.cn/software/gcta/ |
| *plink* 1.9 | <sup>65</sup> | https://www.cog-genomics.org/plink/1.9/ |
| g:Profiler | <sup>66</sup> | https://biit.cs.ut.ee/gprofiler/ |
| ICE CRISPR Analysis Tool | 67 | https://www.synthego.com/products/bioinformatics/crispr-analysis |
| CalmAn 1.9.15 | 34 | https://github.com/flatironinstitute/CalmAn |
| ANTs | 68 | http://stnava.github.io/ANTs/ |
| STITCH | 69 | https://pubmed.ncbi.nlm.nih.gov/27376236/ |
| BEAGLE | 70 | https://faculty.washington.edu/browning/beagle/ |
| cutadapt | 71 | http://code.google.com/p/cutadapt/ |
| BWA | 72 | https://bio-bwa.sourceforge.net/ |
| Python version 3.7 | Python Software Foundation | https://www.python.org |
| Pandas 1.3.4 | 73 | https://pandas.pydata.org/ |
| Pingouin 0.5.2 | 74 | https://pingouin-stats.org/build/html/index.html |
| Matplotlib 3.4.3 | 75 | https://matplotlib.org/ |
| R 4.2.0 | 76 | https://www.r-project.org/ |
| Igraph 0.10.6 | 77 | https://igraph.org/ |
| Other |  |  |
| N.A. |  |  |

### CONTACT FOR REAGENT AND RESOURCE SHARING

Further information and requests for resources and reagents should be directed to and will be fulfilled by the Lead Contact, Su Guo.

### EXPERIMENTAL MODEL AND SUBJECT DETAILS

#### Zebrafish husbandry

All procedures were approved by the University of California, San Francisco Institutional Animal Care and Use Committee. Embryonic, larval, and adult zebrafish were used in this study. Zebrafish were raised at 27-28.5^0^C on a 14/10-hour light/dark cycle. Embryonic and larval zebrafish were raised in blue egg water (0.12 g of CaSO_4_, 0.2 g of Instant Ocean Salts, 30 μl of methylene blue in 1 L of H_2_O) in 90 mm Petri dishes at a density of 40 or less. All assays were conducted on larvae from 6-9 dpf. At this developmental stage, the sex of the organism is not yet determined.

#### Zebrafish GWAS population development

The EKkwill Wildtype (EKW) zebrafish strain with a high level of genetic diversity was used for our GWAS study. The population was derived from 100 pairs of founders from Ekkwill Breeders (Ruskin, FL) using mass breeding. A breeder panel consisting of 142 females and 162 males were used for pairwise mating. A total of 192 larvae derived from all successful crosses were produced for the weekly behavior test (**Figure 2**). Crosses were rotated to maximize genetic diversity. After excluding fish with low quality of phenotypic and genotypic data, 5,759 larvae from 501 distinct pairs of breeding were used for the GWAS analysis.

#### Zebrafish transgenic lines and mutants

The transgenic line *Tg[HuC-H2B-GCaMP6s]* of the AB background and carrying mutations in the pigment synthesis genes (*nacre* and *roy*) were used in this study. Mutants in protein-coding genes and regulatory elements were generated by co-injection of three guide RNAs (gRNAs) and Cas9 protein pre-mixed to form ribonucleoprotein complexes (RNPs) EKW embryos. The resulting mosaic larvae were subjected to behavioral testing followed by genotyping via Sanger sequencing. A subset of RNP-injected larvae were also raised to adults for establishing mutant lines and behavioral testing.

#### Selective breeding of avoidance-inclined SDA and approach-inclined VDA

To develop zebrafish lines with enhanced SDA or VDA behavior, we initiated a selective breeding program (**Figure S7**) with six pairs of SDA and six pairs of VDA individuals (S1) identified from the founder EKW population (S0). The S2-SDAs selected from different S1-SDA families were intercrossed to avoid inbreeding. The VDA lineage was propagated similarly towards the other extreme end of the behavioral spectrum. The two lineages were compared for the seven behavioral indices over the generations (**Figure 7** and **S7**) to verify the cumulative effects of selection. Three individuals from the current generation (S5) of SDA and VDA were stained with HCR *in situ* for *nos1* and *vglut2.2* spatial expression comparison following behavioral selection.

## METHOD DETAILS

### Behavior

#### Light dark preference behavior

##### Behavioral assay

The light-dark preference behavior was performed as previously described ^29^. Fish were raised as a group of 40 in a petri dish on a blue pad at 28.5^0^C with 14-10 hours light/dark cycle and those with normal morphology were individualized in 12-well plates at 5 dpf. At 6 dpf, fish were moved to the behavior room for one hour acclimation followed by two 8-min trials separated by 2.5 hours. Two more trials were carried out at 7 dpf following the same time frame. The behavioral arena is a 4 cm x 4 cm x 1 cm chamber wrapped with opaque tap on the outside wall. The bottom is aligned on top of clear and opaque black acrylic stripes such that the light underneath the acrylic stripes only pass through the clear half of the chamber, thereby generating a half light-half dark compartment. Free swimming behavior was recorded using an infrared filter (Acrylite IR acrylic 11460) fitted Panasonic CCD camera placed above the chamber. Sixteen chambers were recorded simultaneously with the Noldus MPEG recorder 2.1 and stored in MPEG4 format for further analysis with the Noldus Ethovision XT software.

#### Data processing

To characterize the behavior, we used a light dark choice index (LDCI) defined from previous studies that indicates the time a fish spent in the dark zone relative to the light zone. We define those with mean LDCI from 4 trials below the 10-percentile as larvae with strong dark avoidance (SDA) and above the 90-percentile as larvae with varied dark avoidance (VDA). In addition, six other behavioral indices were computed to measure the light-dark preference and locomotor behavior. We computed each index for every trial and its mean across the four trials.

Consequently, we prepared a raw phenotypic matrix that quantifies an individual’s behavior in each row with a total of 35 indices. After regressing out confounding effects arising from each chamber and each week and removing outliers, the phenotypic data was quantile-normalized to satisfy the normality assumption of the GWAS model ^36 39^. For QTL mapping of the indices measured in the later trial, the measurement in the earlier trial within the same day was used as a covariable.

#### Preference behavior for visual cues that mimic food-like particles or predator eye ***spots***

##### Behavioral assay

We tested preference behavior of 9 dpf SDA and VDA individuals for visual cues mimicking either food-like particles or predator eye spots. Four pairs of large black dots (1.2 mm diameter each) resembling four pairs of eye spots were used as a predatory-like visual cue. Forty-eight small black dots (0.5 mm diameter) that occupy the same areas of blackness were used to resemble food-like particles. Visual cues were attached on one inside wall of a transparent behavioral chamber with the same dimension as the light-dark preference chamber. Every experiment was carried out in four 8-min trials on the same day separated by one hour. A control experiment was performed without presenting any visual cue.

##### Data processing

To quantify the responses towards the two visual cues, we divided the behavioral chamber in four virtual zones of equal area and estimated the time fish spent in the zone closest to the visual cue (i.e., zone 1). A two-sample t-test was used to compare zone 1 duration between SDA and VDA presented with the same type of visual cues. Within each group, individuals tested with different visual cues were compared to the control group using the same statistical method.

### In vivo two-photon brain-wide neural activity imaging and data analysis

#### In vivo brain-wide calcium imaging

SDA and VDA larval zebrafish were selected by a light-dark preference behavioral test using the transgenic line *Tg[HuC-H2B-GCaMP6s]* of the AB background and carrying mutations in the pigment synthesis genes (*nacre* and *roy*). At 8 dpf, larva was muscle paralyzed with mivacurium chloride (2mg/ml, final concentration) for 5 min and embedded in 2% low melting agarose on 35 mm Petri dish and covered with 4 ml E3 medium (5 mM NaCl, 0.17 mM KCl, 0.33 mM CaCl_2_, 0.33 mM MgSO_4_). Two-photon Ca^2+^ imaging was performed with a two-photon microscope built based on Thorlabs’s Bergamo series with a resonant scanner. For brain-wide live imaging at 920 nm, we used a 16x objective (0.8 NA; Nikon) covering 1024 x 512 microns field of view at 1.1x zoom. Imaging was carried out at 1 Hz with 1024 x 512 pixel (0.977 microns/pixel). Each whole-brain volume was covered in 28 z-planes separated by 12-μm. Prior to functional imaging, a structural stack was obtained in 168 z-planes with 2-μm spacing.

#### Volume registration of image stacks

All volumes of live brain imaging were registered to Elavl3-H2BRFP stack from the Z-Brain Atlas ^35^. The Z-stack image files were saved in .nrrd format. Registration was performed on the UCSF Wynton local computing cluster by running the following ANTs command antsRegistrationSyN.sh-d 3 -f Z-brain_Elavl3-H2BRFP-1.nii -m $[input].nii -n 8 -o $[output] -t s. The resultant warp and affine coefficient matrices was used for image stack transformation with ANTs command antsApplyTransforms -d 3 -v 0 -n WelchWindowedSinc -I [$input].nii -r Z-brain_Elavl3-H2BRFP-1.nii -o $[output].nii.gz -t $[output]1Warp.nii.gz -t $[output]0GenericAffine.mat. The transformed stack was compared with Elavl3-H2BRFP in Image J.

#### Neuronal activity data processing

The brain-wide Ca^2+^ imaging data were processed with the CaImAn pipeline ^34^ using the parameters from a previous study ^18^. Output files carried extracted regions of interest (ROIs) with the coordinates and df/f (GCaMP fluorescence signal intensity) trace over 900 time points (1s/point). Output files were further processed with a custom written script to assign each ROI to Z-Brain anatomical masks. ROIs outside the brain were excluded as false positive detections ROIs assigned to ‘Ganglia -’, ‘Eyes -’ and ‘Spinal Cord -’ masks were also eliminated in further analysis. An average of 39k ROIs per individual were detected across the 16 fish. A single-neuron activity is measured as the variance of its df/f across the 900 s. Mean variance across neurons assigned to the same mask is used as an estimate of the mask activity. Masks with 0 neuron assigned in any fish were excluded in further analysis.

#### Statistical analysis of neuronal activity data

Since the whole brain activity and the activity of the four broad regions (“Diencephalon –“, “Mesencephalon –“, “Rhombencephalon –“ and “Telencephalon –“) showed no significant differences between SDA and VDA, we compared the activity of all other subregions between the two groups. We used median of ratios method to normalize the activity of each mask as it combines the inter- and intra-sample variations to provide a sample-specific adjustment factor. Specifically, a pseudo-reference activity was created for each mask by taking the mask geometric mean across the samples. A ratio of the mask activity over the pseudo-reference activity was computed for each mask in a sample. Finally, the median of the ratios across the masks within a sample was used to divide each mask activity of the sample. For statistical analysis, the normalized data were first tested with one-way ANCOVA to generate an experimental *p*-value for each mask. The activity of the relevant broad region was used as the covariable. To control for false positive rate, we permuted the dataset with one-way ANCOVA for 1,500 rounds. The phenotypic category (i.e., SDA and VDA) was shuffled in each permutation round without repetition to generate an empirical *p*-value null distribution for adjusting the experimental *p*-value, leading to a detection of mask set 1 that distinguished SDA from VDA. For masks showing minimal correlation with the covariable, we performed a one-way ANOVA permutation test to discover a second mask set. Masks that showed neuronal activity differences between SDA and VDA in either set was considered as significantly different between the two groups.

### Genotyping

#### DNA isolation

DNA was isolated from whole larvae of 6 dpf old or from fin clips obtained from an adult zebrafish using DNAdvance kit (Beckman Coulter, Brea, CA) in 96-sample batches according to manufacturer’s protocol. Briefly, tissue samples were placed in 96-well plates and incubated with proteinase K at 55°C overnight using Thermomixer module, then DNA was bound to SPRI beads, washed with ethanol and eluted. DNA concentration was measured using Nanodrop (Thermo Fisher Scientific, Waltham, MA).

#### Sequencing

For the low-coverage sequencing, we used Twist 96-Plex Library Preparation kit (TWIST, San Francisco, CA) library preparation method, which is based on random priming. Libraries were prepared from 100 ng DNA in batches of 96 samples, then batches of 960 samples were loaded onto a single lane of Illumina NovaSeq 6000 and sequenced as PE150. Reads were demultiplexed with Fgbio software (Fulcrum Genomics, fulcrumgenomics.github.io/fgbio/), trimmed with cutadapt software and then aligned to zebrafish genome assembly GRCz11 using BWA. Duplicates were marked using Picard MarkDuplicates function. The mean coverage was 0.5x. Samples that were used for reference panel and for concordance check were deep-sequenced. The libraries were prepared using KAPA Hyper method and then sequenced on Illumina NovaSeq 6000 to generate ∼30x coverage (range 20x – 55x). Sequencing reads were trimmed with cutadapt software. The reads were aligned to zebrafish genome assembly GRCz11 using BWA. Duplicates are marked using Picard MarkDuplicates function.

#### Variant calling

Samtools were used to process the 94 deeply sequenced (30 x coverage) F_0_ breeders. The 8 deeply sequenced control study F_1_ samples were processed using STITCH ^69^, the same way as for other study samples sequenced by lcWGS (0.5 x coverage), now that the positions of a list of variants were already obtained from the 94 breeders. For STITCH, we used the parameters K = 60 and nGen = 2. We used niterations = 1 parameter because we had 94 F0 breeder sequenced at 30x and used their genotypes as a reference set. Then imputation to the reference was performed using BEAGLE (ne = 106).

Since STITCH would produce similar results with about the same precision no matter if all the samples were processed together or in small sets, we performed imputation for 2 sets of individuals (3,373 and 2,594), and then run BEAGLE again on two sets combined. Samples that had >25% missing genotypes after STITCH were excluded from consideration. SNPs that were missing in >15% of samples after STITCH were excluded from consideration. The accuracy of the imputation was measured by using a truth set containing 7 F1 individuals sequenced at 30x. These individuals were also sequenced at 8x using Twist library prep method, to generate 50 pseudo-samples to reflect the distribution of the number of reads across all shallow-sequenced zebrafish samples. The discordance rate that was measured for these pseudo-samples, in comparison to the 7 deep-sequenced samples, was 1.67%.

#### Genotype quality control

Filtering raw genotypic data yields a population of 5759 individuals with 20.49M SNPs for further quality control. Extremely rare variants (MAF<1%), variants with low STITCH info score (<0.75) and variants violating Hardy-Weinberg equilibrium (*p*-value > 10^e-6^) were eliminated. After LD prune (r^2^>0.95), 4.31M SNPs were used for GWAS analysis.

### Pre-GWAS analysis

#### SNP heritability

The SNP heritability, which measures the proportion of variance attributable to all SNPs, was estimated using GCTA-GREML analysis (**Figure S3A**).

#### Phenotypic and genetic correlation between traits

Phenotypic correlations between the seven mean behavioral indices were determined using the *Pearson* correlation coefficient. Genetic correlations were calculated using bivariate genome-based restricted maximum likelihood analysis (GREML) implemented by the –reml-bivar function in GCTA (**Figure S3B**).

#### LD decay

The linkage disequilibrium decay of the EKW population was determined by computing pairwise LD statistic *r^2^* ^65^ using the --r2 function in *plink*. Common SNP pairs (MAF>20%) with similar frequencies (differ by less than 5%) and separated with an interval ranging from 0-100kb to 4.9-5Mb were collected. A mean LD estimate for each interval was obtained to produce the LD decay plot (**Figure S2G**).

#### Population stratification test

The --pca and --ibc function in *plink*^10^ was implemented to perform PCA analysis and to estimate individual inbreeding coefficient, respectively (**Figure S2A-D**).

### GWAS for mapping QTLs

We implemented a linear mixed model (LMM) with the genotypic and phenotypic data of 5,759 larvae using GCTA ^64^. GCTA estimates both fixed and random effects for each marker, which can deflate test statistic and reduce the power to detect a QTL. This issue, termed as proximal contamination, was addressed by using the leave-one-chromosome-out (LOCO) approach that tests a marker with a genetic relationship matrix (GRM) constructed by eliminating markers from the same chromosome ^41^. We used a permutation-based approach to calculate the genome-wide significance threshold for *p*-values calculated in GCTA (**Figure S3C**). A null distribution of p-values was generated by mapping QTLs in 1,200 randomly permuted data sets and the 95 percentile of this distribution was taken for thresholding. We applied the same threshold to all behavioral indices since they were quantile normalized. To establish independency of significant SNPs that are nearby one another, we performed a stepwise regression approach that re-exam the SNP effects conditioned on the effect of the SNP with lowest *p*-value. This process was repeated (including all previously significant SNPs as covariates) until no more QTLs were detected on a given chromosome. Peak SNPs passing the independency test were plotted in **Figure 3A** as surrogates for the QTL regions. Coordinates of the furthest marker showing a high LD (r^2^ >= 0.6) with the peak SNPs were used to define the intervals of each QTL.

### Bioinformatics analysis of GWAS candidate genes

We collected protein-coding genes from all the GWAS QTLs as candidates for bioinformatics analysis. For small QTLs harboring less than three genes, we loosened the LD stringency by using r^2^ = 0.5 to include more candidate genes since causal effects can be exerted from a distance larger than the size of QTLs ^78^. Of the resulted gene set, we focused on 338 candidates that have human orthologues comprehensively annotated in the Gene Ontology database. We performed an enrichment analysis with the GO terms of these human orthologues using a web server g:profiler to identify significantly enriched GO terms (adjusted *p*-value < 0.05). For each enriched GO term, we downloaded its directed acyclic graph (DAG), the original hierarchy in GO database, and reformed it using an R package igraph followed by term re-ranking with the sugiyama algorithm. Starting from the end nodes, the lineage of each term was determined by the longest path. For paths of equal length, the node with better description was manually selected. The output lineage was represented in a sunburst plot using a python package plotly (https://plotly.com/). Since the most enriched GO terms were related to synaptic transmission, we then cross-referenced our human gene set with the human GWAS catalog database ^42^ for overlapped genes associated with three neural related categories including psychiatric disease, mental processes and neurodegenerative disease. The psychiatric disease with the most overlapped genes was further partitioned into five common diseases namely schizophrenia (SCZ), unipolar depression (UD), attention deficit hyperactivity disorder (ADHD), autism spectrum disorder (ASD) and substance abuse (SBA).

### Discovery and bioinformatics analysis of GWAS significant variants

We retrieved all SNPs to rerun a GWAS analysis in each QTL. The enlarged set of significant SNPs were queried for its putative regulatory effects using the DANIO-CODE, a central repository of zebrafish developmental functional genomic data ^43^. We interpreted potential functional alteration caused by these significant SNPs using Ensembl Variant Effect Predictor (VEP) ^79^. We downloaded multiple alignment sequence files across eight species (including zebrafish, medaka, stickleback, tetraodon, fugu, frog, mouse, and human) from the UCSC Genome Browser and analyzed the alignments with Biopython to discover conserved elements that contain our GWAS significant SNP variants. Regulatory function of the elements conserved between zebrafish and human were further explored using RegulomeDB ^44^.

### Multiplexed Crispr-Cas9 mediated genome editing

We used a multiplexed CRISPR-Cas9 mediated knockout method in F0 animals as previously described ^80^. We designed three gRNAs targeting distinct 5’-most exons of each gene. For genes with multiple transcript variants, exons conserved across all the transcripts were targeted. We queried CRISPRscan, ChopChop, and IDT Guide RNA design checker tool for gRNA design and those predicted with high activity score and low off-target rate in all three databases were selected. The three designed gRNAs were microinjected as multiplexed RNPs into the yolk of zebrafish embryos at one-cell stage to generate mosaic F0 animals. At 6 dpf and 7 dpf, larvae with normal morphology were behaviorally tested. After that, DNA of five samples from each group were individually PCR amplified for a 200-300 bp fragment containing each sgRNA target site. Bulked PCR products were used for sanger sequencing followed by knock-out efficacy assessment using the ICE Analysis Tool (Synthego Performance Analysis, ICE Analysis. 2019. v3.0. Synthego).

### HCR *in situ* hybridization

To investigate the effect of knocking out the putative regulatory element (*re-nos1*) on *nos1* expression, we used six larval zebrafish homozygous for a ∼ 300 bp deletion in the conserved sequence element (encompassing a GWAS significant variant in this QTL). Their head tissues were processed for HCR *in situ* and results were compared with four siblings without the large deletion. To determine the effects of natural genetic variations (not limited to this *re-nos1* element) on *nos1* expression, we applied HCR *in situ* to three S5-SDA and three S5-VDA from our selective breeding program.

A set of DNA antisense oligonucleotide pairs carrying split B3 initiation sequence were designed by Molecular Instruments (Los Angeles, California, USA) to tile across the length of the *nos1* mRNA transcript. Another set of probes were designed to anneal with B1 initiation sequence to hybridize to the *vlgut2.2* mRNA for registration and as a control. Dye-conjugated hairpins (B1-546 and B3-488) and other buffers were purchased from the same company. Larval zebrafish were stained individually in a 250 ul PCR tube throughout the entire procedure according to a modified protocol of the “HCR v3.0 protocol for whole-mount zebrafish embryos and larvae” provided by Molecular Instruments ^45^. After behavioral test, 7 dpf fish were fixed with ice-cold 4% PFA/PBS overnight at 4°C with gentle shaking. After washing 3 x 5 min with 1x PBST, fish were treated with a bleach solution (0.18 M KOH and 3% H_2_O_2_) for 10 min followed by 1% PBTriton washing (3 x 5 min) to remove pigments. Bleached fish underwent a short 10 min treatment with ice-cold 100% Methanol at −20°C to dehydrate and permeabilize the tissue samples. Next, rehydration was performed by serial washing of 50% MeOH/50% PBST and 25% MeOH/75% PBST for 5 min each and finally 5 x 5 min in 1x PBST. For probe hybridization, fish were pre-hybridized with pre-warmed hybridization buffer for 30 min at 37°C. Probe solution was prepared by mixing 1.6 pmol of each HCR probe set with 100 μl of hybridization buffer and was used to replace the hybridization buffer for a 16-hour incubation at 37°C. To remove excess probes, fish were washed 4 x 15 min with 100 μl of pre-warmed probe wash buffer at 37°C. Subsequently, larvae were washed 3 x 5 min with 5x SSCT (5x sodium chloride sodium citrate + 0.1% Tween-20) buffer at room temperature. Next, preamplification was performed by incubating each sample in 50 μl of amplification buffer for 30 min at room temperature. Meanwhile, 7.2 pmol of hairpin h1 and 7.2 pmol of hairpin h2 were heated to 95 °C for 2 min then cooled down to room temperature in a dark environment. For hairpin amplification, the preamplification buffer was replaced with a hairpin solution made by adding hairpins to 50 ul amplification buffer for a final concentration of 120 nM. Samples were incubated in the hairpin solution for 16 hours in the dark at room temperature. Fish were washed the next day 3 x 20 min using 5x SSCT at room temperature to remove excess hairpins and were mounted in 1.5% low-melting-point agarose and covered in di-H_2_O and imaged within 5 hours using Leica Stellaris 8 Tau STED confocal scanning microscope (inverted) equipped with 20x water immersion objective. A two-channel z-stack file composing 2 tiles were taken for each fish to produce a final image of 971×517 pixel (883 x 470 μm, 2.5 μm in z). Both channels were imaged in bidirectional resonant scanning mode (2× line average) without crosstalk.

### HCR *in situ* image data processing

To construct a template with *vglut2.2* HCR hybridization signal arranged in Z-brain atlas (hereafter referred as Z-brain HCR-*vglut2.2*), we aligned mapzebrain ^81^ H2B-GCaMP2 to Z-brain H2B-RFP and apply the transformation to mapzebrain HCR-*vglut2.2* (**Figure S6**). All HCR Z-stacks were registered to the Z-brain HCR-*vglut2.2* using the same method as described for *in vivo* calcium imaging. After registration, the 138 slices of the *nos1* channel were preprocessed individually using a four-step procedure. First, slices were convolved with an isotropic Gaussian blur (image J, standard deviation sigma of the Gaussian) to remove noise. To eliminate pixel variations arising from background, a local background subtraction was applied with the “rolling ball” algorithm (image J). Next, pixel intensities were normalized within each slice to eliminate imaging variations due to slice depth. Finally, pixels were filtered with the 90^th^ percentile and the retained pixels were assigned to the Z-brain atlas anatomical mask. The number of assigned pixels was used to measure the *nos1* expression level in each anatomical mask. The same processing pipeline was applied to *vglut2.2* HCR imaging data.

### Statistical analysis for spatial gene expression

We performed two comparisons of HCR *in situ* data (i.e., *re-nos1* KO vs sib, S5-SDA vs S5-VDA) with the same statistical analysis pipeline used for *in vivo* calcium imaging data analysis except that the maximum rounds of permutation were determined according to the sample size of each group (210 for *re-nos1* KO vs sib, 20 for S5-SDA vs S5-VDA). To verify if the detected *nos1* spatial expression differences are gene specific, we performed the same analytical process with *vglut2.2* HCR expression data.

### Correlation analysis of *nos1* and *vglut2.2* spatial expression

Masks that are anatomically closely located or functionally related might share similarities in gene expression. To that end, we assessed the expression correlation between the masks for each gene by using 30 larvae with varied light-dark preference. Only masks with non-zero expression in all 30 larvae were analyzed. In addition to analyzing *nos1-nos1* and *vglut2.2-vglut2.2* correlation, we investigated the correlation between the two genes for all pairs of masks with the co-expression of the two genes. We performed pairwise *Pearson* correlation analysis followed by false discovery rate control with *Benjamini–Hochberg* procedure.

## SUPPLEMENTARY FIGURES

**Figure S1.**
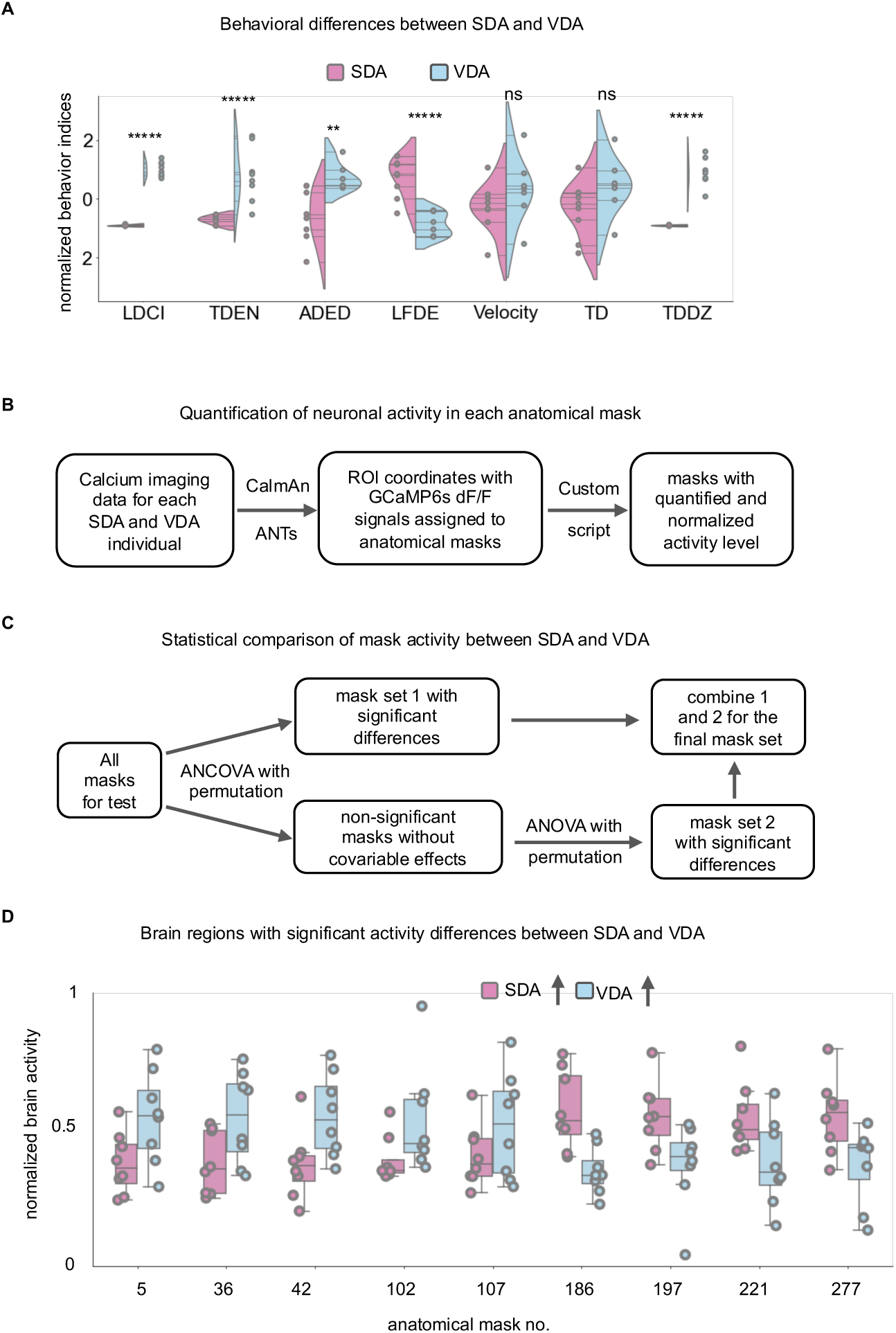
Behavioral and neural activity analyses of approach-inclined VDAs and avoidance-inclined SDAs. Related to Figure 1. **A**. Behavioral indices in the light-dark preference paradigm of SDAs and VDAs selected for brain activity analysis. One-way ANOVA, *p*-values are adjusted with *Benjamini-Hochberg* false discovery rate correction. \**p*<0.05, \*\**p*<0.01, \*\*\**p*<0.001, \*\*\*\**p*<0.0001, \*\*\*\*\**p*<0.00001. n.s. not significant, n=8 per group. LDCI (Light-Dark Choice Index), TDEN (Total Dark zone Entry Number), ADED (Average Dark zone Entry Duration), LFDE (Latency to First Dark zone Entry), TD (Total Distance traveled), and TDDZ (Total Distance traveled in the Dark Zone). **B**. Pipeline of brain-wide neural activity data processing. Calcium imaging data were processed through CaImAn and the coordinates of identified ROIs (individual neuronal nuclei) were registered to the Z-brain Elval3-H2BRFP using ANTs toolkit. **C.** Statistical workflow for detecting brain regions with significant neural activity differences between SDAs and VDAs. One-way ANOVA and ANCOVA permutation tests, significance is indicated if adjusted *p*-value < 0.05. **D.** Box plots show nine anatomical masks with significant neural activity differences between SDAs and VDAs. Masks are numbered according to the Z-brain atlas.

**Figure S2.**
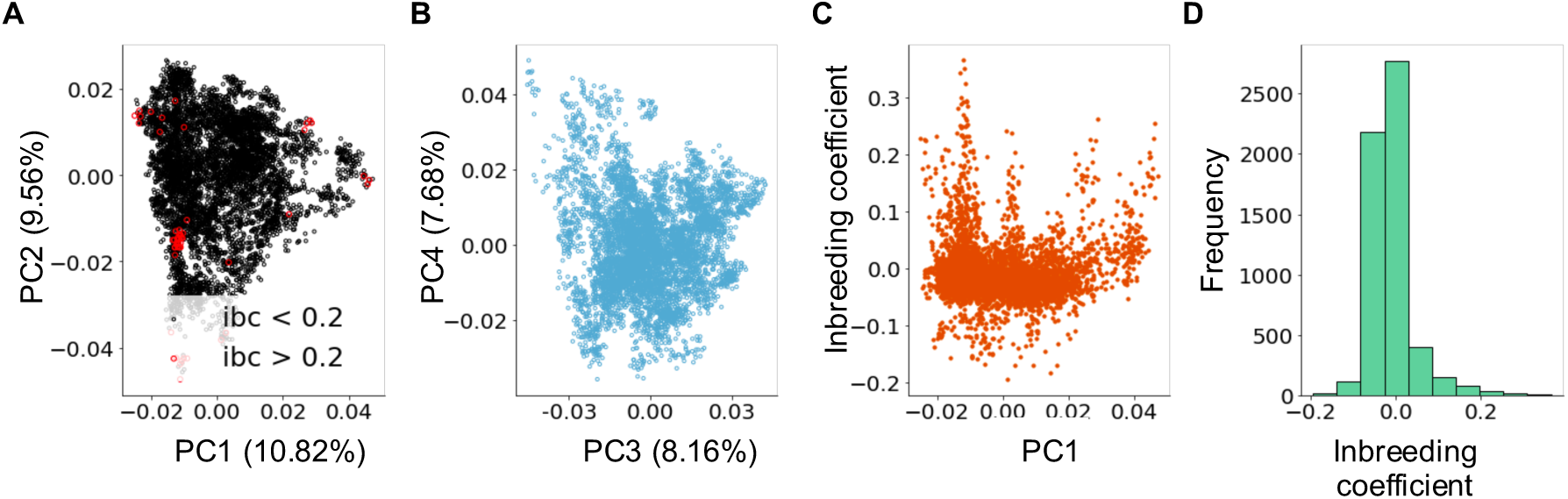
Population structure of EKW zebrafish used for GWAS. Related to Figure 2. **A and B.** Top 4 principal components (PC1 and PC2 in **A**, PC3 and PC4 in **B**) of genotype dosage matrix. Each dot is an individual subject. Dots in **A** is colored based on its inbreeding coefficient (*ibc* < 0.2, black; *ibc* > 0.2, red). **C**. Inbreeding coefficient shows no correlation with the first principal component. **D**. Distribution of inbreeding coefficient. Most individuals have inbreeding coefficients near 0, indicating our mating scheme has maintained an outbred GWAS population.

**Figure S3.**
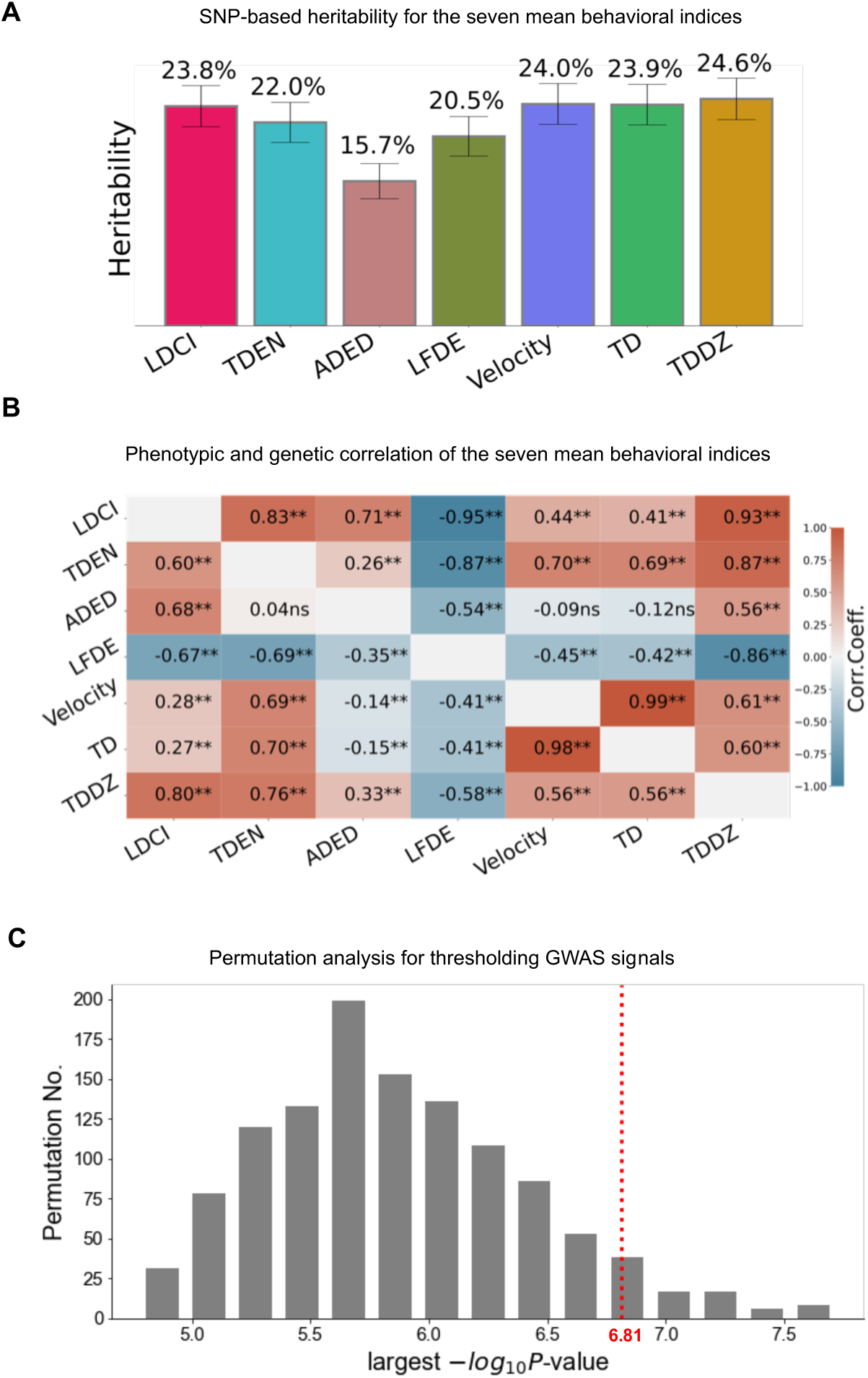
Analyses of GWAS-related data. Related to Figure 3. **A**. SNP-based heritability estimates (percent trait variance explained by 4.31 million GWAS SNPs) for the eight mean behavioral indices (mean ± s.e.m). LDCI (Light-Dark Choice Index), TDEN (Total Dark zone Entry Number), ADED (Average Dark zone Entry Duration), LFDE (Latency to First Dark zone Entry), TD (Total Distance traveled), and TDDZ (Total Distance traveled in the Dark Zone). **B**. Correlation at the genetic (upper part) and phenotypic levels (lower part) among the 7 behavioral indices. The genetic correlation is estimated by applying the GCTA –reml function to the genetic relationship matrix (grm). The phenotypic correlation is estimated with *Pearson* correlation. Correlation coefficient is indicated inside squares with significance symbolized (null hypothesis: correlation coefficient = 0, \**p* < 0.05, \*\**p* < 0.01. n.s. not significant. *Benjamini– Hochberg* false discovery rate correction for correlation test across all indices pairs. **C**. Results of permutation analysis for QTL mapping. The original dataset is permuted for 1,500 rounds to generate an empirical distribution of the minimum *p*-value from each permutation. The minimum *p*-value in each permuted dataset is the largest −log_10_*P*-value from the 4.31 million SNPs tested for association. This permutation test is meant to simulate the distribution of *p*-values under the null hypothesis. The 95^th^ percentile of this distribution, which is approximately 6.81, is depicted here by the dotted red line.

**Figure S4.**
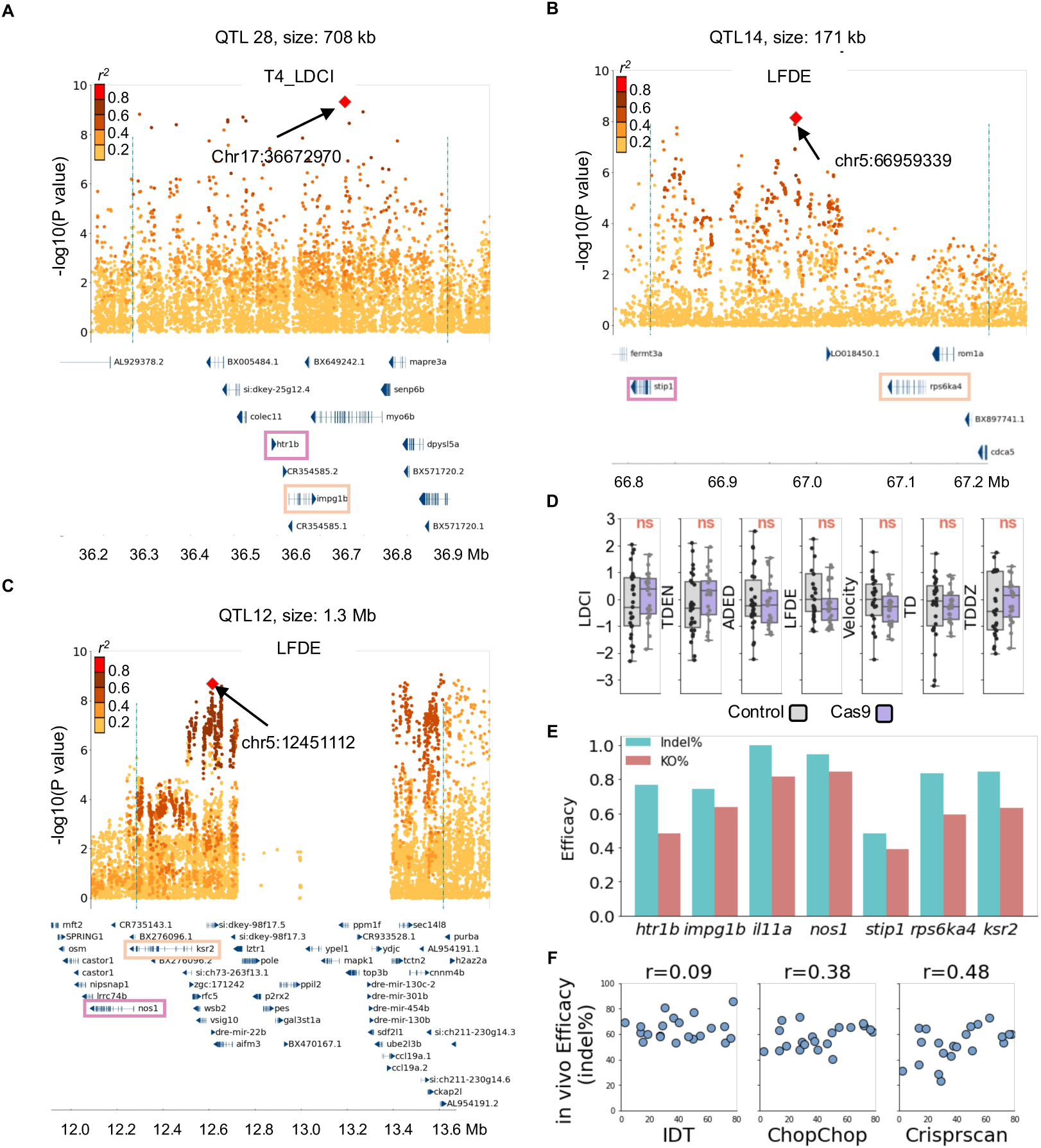
Functional validation of GWAS candidate genes. Related to Figure 4. **A, B,** and **C**. Locus Zoom plots for QTLs containing the three casual genes. The causal gene *htr1b* (serotonin receptor 1b) is in QTL #28 that spans 708 kb on chr. 17. The causal gene *stip1* (stress-induced phosphoprotein 1) is in QTL #14 that spans 171 kb on chr. 5 (B). QTL #12 (1.3 Mb) is on a different region of chr. 5 (C) and harbors the third casual gene *nos1* (neuronal nitric oxide synthase 1). **D.** Embryos injected with Cas9 only are indifferent from the sibling control in all seven behavioral indices. Two-sample t-test. n.s. not significant, n=27, *p*-values are adjusted by *Benjamini–Hochberg* false discovery rate correction for test across all indices. **E**. *In vivo* efficacy measurement of multiplexed *Crispr-Cas9* gene editing method. After behavioral testing, DNA from five RNP injected larvae is isolated for a 200-300bp PCR product centered at a gRNA PAM site. The PCR products are pooled for sanger sequencing together with a control PCR product amplified from the sibling DNA. The sequence file is processed by ICE to estimate the KO rate of each gRNA. A joint KO rate across the three gRNAs designed for the same gene is considered as the KO rate of the targeted gene (red). Similarly, a joint rate of generating an indel in the gene is computed using the indel rate of each individual gRNA. **F.** Correlation between *in vivo* and *in silico* estimate of the tested gRNA efficacy. The indel rate of each gRNA tested with sanger sequencing and ICE is compared with its predicted efficacy in three sgRNA design software.

**Figure S5.**
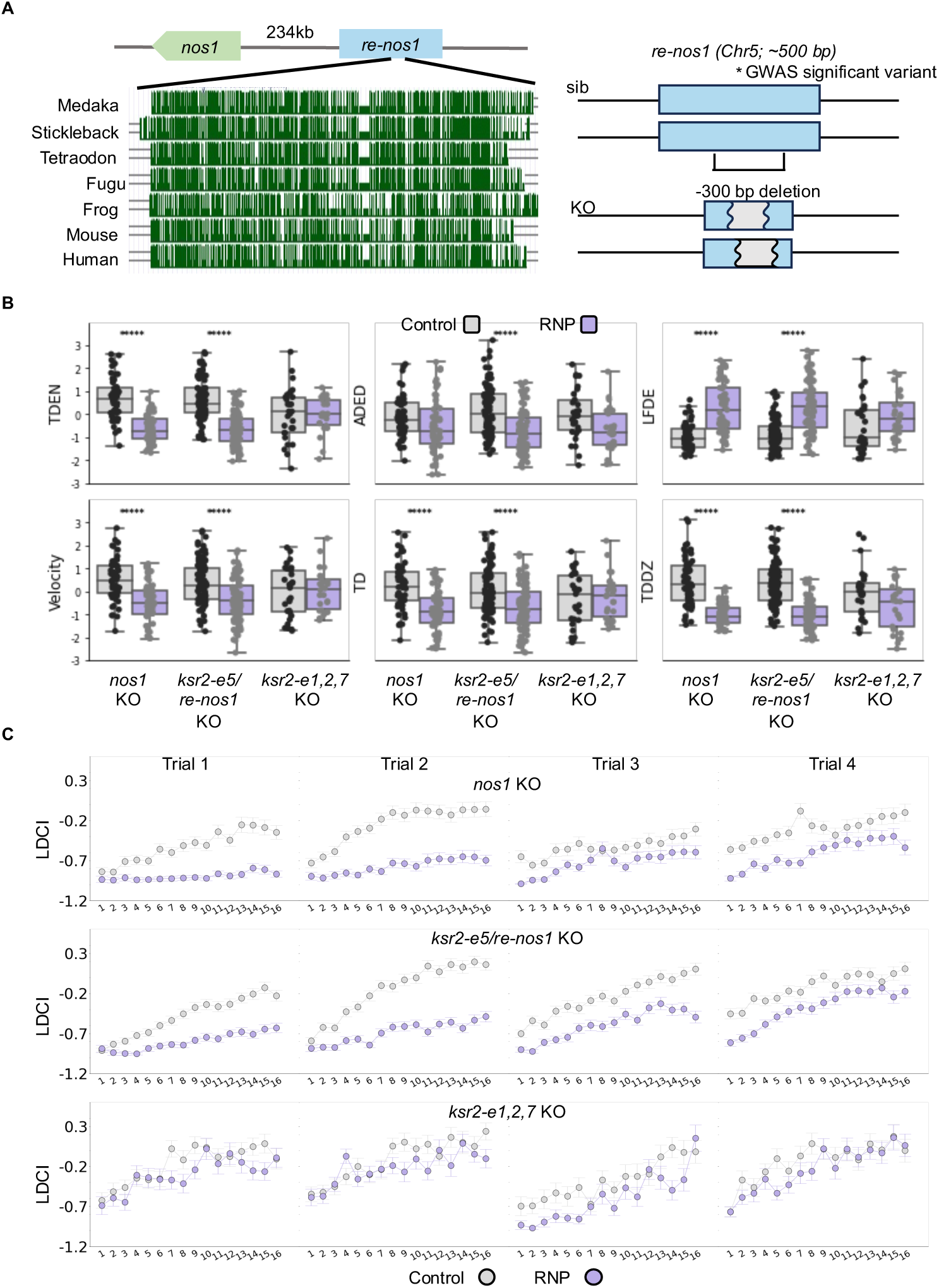
Behavioral effects of knocking out *nos1*, *ksr2-e5/re-nos1*, and *ksr2-e1,2,7*. Related to Figure 6. **A**. Schematic shows the conservation of *re-nos1/ksr2-e5* and the nature of the KO. A ∼300 bp deletion is generated in the targeted region using the multiplexed genome editing. The deletion is verified by sanger sequencing. **B**. Quantification of additional behavioral indices in the *nos1* KO, *ksr2-e5/re-nos1 KO*, and *ksr2-e1,2,7* KO. TDEN (Total Dark zone Entry Number), ADED (Average Dark zone Entry Duration), LFDE (Latency to First Dark zone Entry), TD (Total Distance traveled), and TDDZ (Total Distance traveled in the Dark Zone). **C**. High-resolution behavioral analyses show that the behavioral dynamics are similarly altered in the *ksr2-e5/re*-*nos1* KO and *nos1* KO. Each 8-min trial is divided into sixteen 30 s time-bins. Each dot is the summary data (mean ± s.e.m) from the RNP injected larvae (purple) and the sibling controls (gray). Such changes of behavioral dynamics were not observed in the *ksr2-e1,2,7* KO.

**Figure S6.**
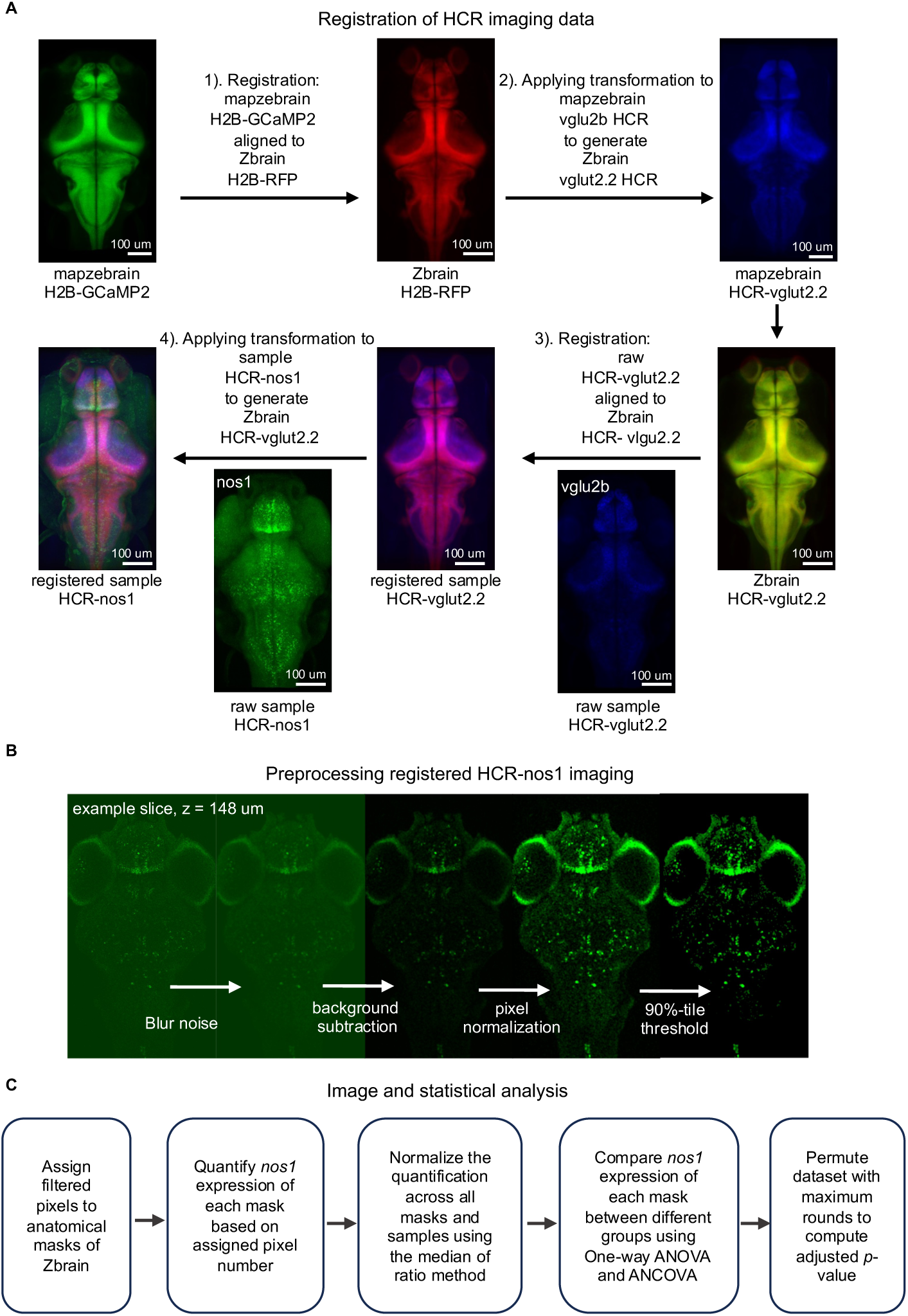
Quantitative spatial gene expression analysis pipeline. Related to Figure 6. **A**. Protocol for registration of the HCR *in situ* imaging data to the Z-brain atlas template. Transformation matrix for registration of mapzebrain template to Z-brain template is constructed by aligning mapzebrain H2B-GCaMP2 to Z-brain H2B-RFP. Applying the transformation to mapzebrain HCR-vglut2.2 results in a Z-brain HCR-vglut2.2 which is then used for the registration of sample HCR imaging data. All the registrations are performed using ANTs toolkit. **B**. An example slice of *nos1* HCR *in situ* data undergoes preprocessing following image registration. The procedure includes four steps to increase the signal to noise ratio. Slices are processed individually using custom Fiji macro and Python script to minimize imaging variations caused by depth. **C**. Workflow of quantification and statistical analysis of *nos1* HCR signals. Filtered pixels are counted equally as 1 regardless of the intensity differences. Mask level intensity is estimated by counting the assigned pixel number. Methods of normalization and statistical test are the same as that for analyzing the brain activity data.

**Figure S7.**
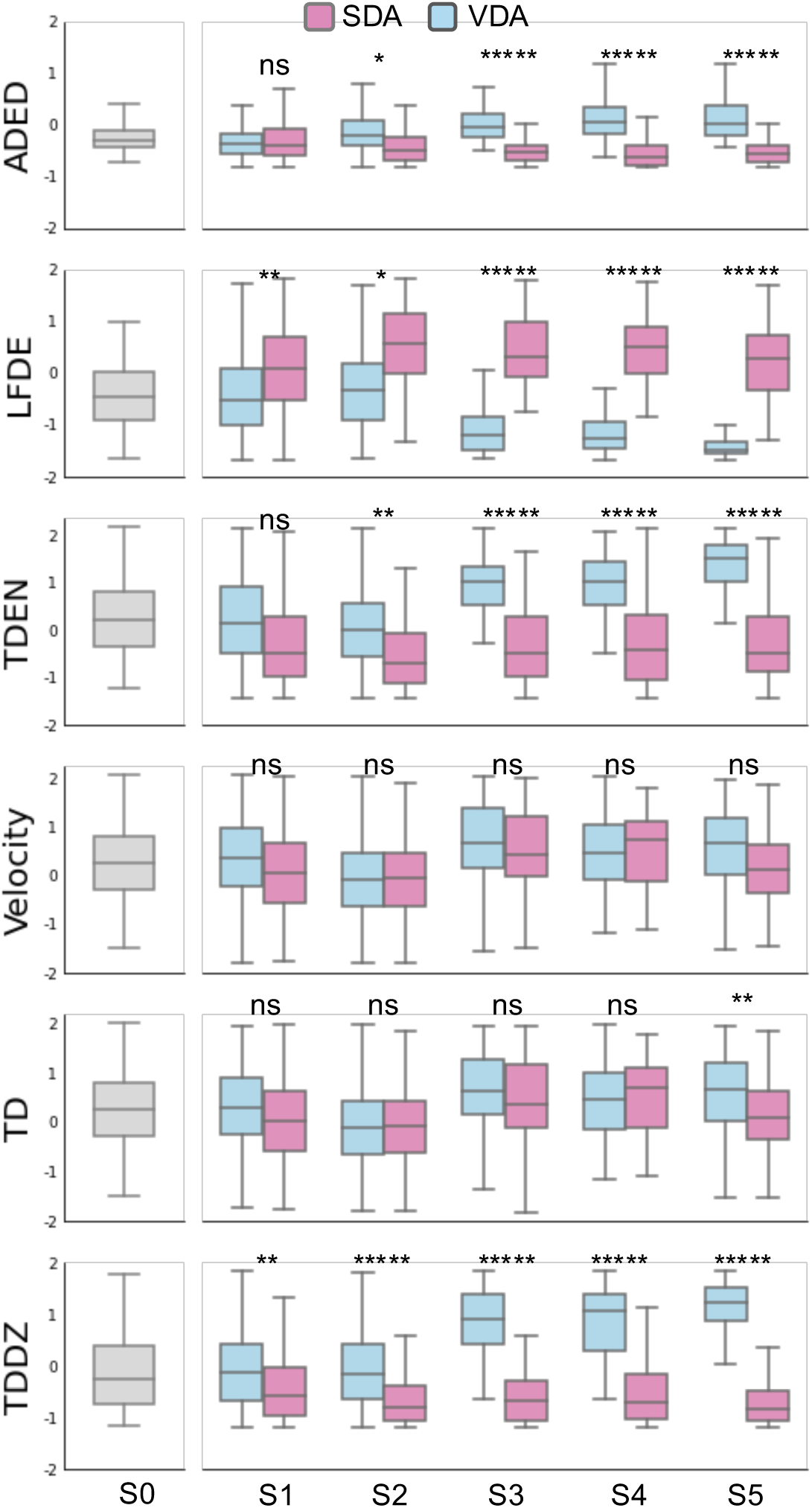
Changes of additional behavioral parameters during selective breeding. Related to Figure 7. In addition to the LDCI, selective breeding takes effect on four other behavioral parameters including ADED (Average Dark zone Entry Duration), LFDE (Latency to First Dark zone Entry), TDEN (Total Dark zone Entry Number) and TDDZ (Total Distance traveled in the Dark Zone). In contrast, velocity was not affected, and the TD (Total Distance traveled) shows a slightly detectable difference in the latest generation. Two-sample t-test, \**p*<0.05, \*\**p*<0.01, \*\*\**p*<0.001, \*\*\*\**p*<0.0001, \*\*\*\*\**p*<0.00001. n.s. not significant, n=35 for each generation. All *p*-values are adjusted with *Benjamini–Hochberg* FDR control across all indices.

